# High-resolution transcriptome analysis reveals a multilayered and dynamic transcriptional architecture in the archaeal virus SSV1

**DOI:** 10.64898/2026.06.17.732947

**Authors:** Wouter Magnus, Maarten Boon, Jorien Poppeliers, Jonathan Abshier, Kenneth Stedman, Rob Lavigne, Eveline Peeters

**Affiliations:** Research Group of Microbiology, Department of Bioengineering Sciences, Vrije Universiteit Brussel, Brussels, Belgium; Laboratory of Gene Technology, Department of Biosystems, KU Leuven, Leuven, Belgium; Center for Life in Extreme Environments, Department of Biology, Portland State University, Portland, Oregon, USA

**Keywords:** Archaea, *Fuselloviridae*, *Saccharolobus solfataricus*, antisense RNA, transcription regulation

## Abstract

*Fuselloviridae* infecting thermoacidophilic *Sulfolobales* are among the best-studied archaeal viruses, with the *Saccharolobus* Spindle Shaped Virus 1 (SSV1)-*Saccharolobus solfataricus* interaction serving as a model system. After infection, SSV1 establishes a chronic infection with new virions continuously budding from the infected cell. UV irradiation triggers virus replication, increased virion production and a tightly orchestrated temporal transcription cycle. However, gene regulatory mechanisms post-UV-induction remain elusive. In this study, we investigated the SSV1 transcriptomic landscape post-UV-induction using high-resolution, long-read ONT-cappable-RNA-sequencing. Through end-to-end sequencing of primary transcripts, we generated a comprehensive transcript map of SSV1, identifying 21 transcription start sites (TSSs), including 12 newly described sites, and 19 termination sites (TTSs), of which 9 are novel. This enabled refinement of transcription units and operon structures across the genome. Five new antisense RNAs were discovered, including one encoded antisense to the open reading frame of a putative viral toxin gene *a291*, which we hypothesize to be a *cis*-encoded regulatory RNA. In addition, spontaneous recombination between two direct repeats in the SSV1 structural genes *vp1* and *vp3* was confirmed. To our knowledge, this is the first implementation of cappable-seq for an archaeal virus, and the first base-resolution analysis of the SSV1 transcriptome. It reveals a previously unappreciated level of transcriptional complexity, illustrating how high-resolution transcriptomics can deepen understanding of viral transcriptome architecture and regulatory networks.

**Importance:** Archaeal viruses remain among the least understood viruses in the virosphere, despite their ecological and evolutionary significance. *Saccharolobus* Spindle Shaped Virus 1 (SSV1) is a key model for the study of virus-host interactions in archaea. Yet, its gene regulatory mechanisms have remained poorly resolved. Here, we provide the first base-pair resolution transcriptome of an archaeal virus, revealing a complex and multi-layered transcriptional architecture. We uncover alternative transcription start and termination sites, as well as antisense transcription, pointing to regulatory mechanisms beyond a simple temporal gene expression program. These findings suggest that archaeal viruses use sophisticated transcriptional and post-transcriptional mechanisms similar to those in bacterial and eukaryotic viruses. By establishing a high-resolution map of the SSV1 transcriptome, this work advances our understanding of archaeal virus biology.

## Introduction

Archaeal viruses remain one of the most enigmatic and understudied components of the global virosphere (1). They are extremely diverse, displaying fascinating morphologies that are not observed among bacterial and eukaryotic viruses, and exhibit extensive variability in genome sequences. Many lineages of archaeal viruses cluster separately from the rest of the virosphere in genomic sequence analyses, making it very difficult to draw evolutionary connections and highlighting the extent of “viral dark matter” that remains to be discovered (2, 3).

*Sulfolobus* spindle-shaped virus 1 (SSV1) was one of the first archaeal viruses to be discovered in 1982 and was first described as a UV-inducible plasmid of its natural host *Saccharolobus shibatae* B12, which was isolated from a volcanic hot spring at Beppu, Japan. *S. shibatae* B12 grows at temperatures above 70°C and pH below 4 (4, 5). SSV1 is capable of infecting *Saccharolobus solfataricus* and establishing a chronic productive infection (6, 7). In the carrier state, its double-stranded DNA (dsDNA) genome is integrated into the host genome at the locus of a tRNA^Arg-CCG^ gene as a linear copy, while a handful of episomal copies remain present in the cytoplasm, and new virions egress from the host through a non-lytic budding mechanism, reminiscent of eukaryotic enveloped viruses (6–10).

SSV1 has been used as a model system for the study of archaeal promoter and terminator elements and proved instrumental in demonstrating that these are homologous to their eukaryotic counterparts (11, 12). Indeed, for multiple SSV1 promoters, promoter elements were identified (8, 11, 13, 14), leading to insights that the archaeal TATA box is a 6-8 base pair (bp) AT-rich sequence, generally displaying the motif TTTAWA(TR), centered around 27 bp upstream of the transcription start site (TSS) (15–17). Directly upstream of the TATA box, a conserved purine-rich Transcription Factor B Recognition Element (BRE) is typically present, displaying the general motif RNWAAW (18). Closer to the TSS, around positions −12 to −1, an AT-rich sequence termed the promoter proximal element (PPE) is found, followed by the initiator element (Inr), which consists of a typical pyrimidine-purine dinucleotide at the TSS (positions −1 to +1) (16, 17).

Increased SSV1 virion production is induced by UV irradiation or by DNA-damaging agents such as mitomycin C (19, 20). Besides a burst of new virions, a temporal transcriptional program has been observed for SSV1 genes upon UV induction, illustrating how transcription regulation might be key to the SSV1 life cycle (11, 13, 14). However, it remains unresolved how the virus manages this regulation at the molecular level. A substantial fraction (12 out of 35) of SSV1 open reading frames (ORFs) has been predicted to encode DNA-binding proteins, which could be involved in this regulation (21). Nevertheless, only one of these proteins, namely F55, has been functionally characterized as a regulator of transcription of the transcripts T5, T6, Tind and Tlys, involved in the switch from carrier state to induced state (8, 22, 23). These first studies mapping SSV1 transcripts, quantifying virus induction and describing the transcription cycle have laid invaluable groundwork for further SSV1 research. However, these analyses were performed using low-resolution technologies such as S1 endonuclease mapping of transcripts, electron microscopic counting of virions, semi-quantitative PCR and DNA microarray analysis. As a result, the precise architecture of the SSV1 transcriptome, including exact transcription boundaries and transcript isoforms, has remained unresolved.

Thanks to substantial technological advancements over the past decades, we can now use high-resolution techniques such as Oxford Nanopore Technologies-based cDNA cappable sequencing (ONT-cappable-seq), to perform in-depth transcriptomics and map the SSV1 transcriptional landscape at base pair resolution (24). This cappable-seq approach is based on the enrichment of primary mRNAs, which contain valuable information on transcription start sites (TSSs), transcription termination sites (TTSs), operon structures and untranslated regions (UTRs), from the RNA pool consisting of mostly processed transcripts. Combining cappable-seq with end-to-end readout of these primary mRNAs using Oxford Nanopore enables direct delineation of full-length transcription units and regulatory elements, providing a complete blueprint of the virus transcriptome.

In this study, UV induction of SSV1 genome replication is revisited and quantified using real-time quantitative PCR (qPCR), providing a well-defined experimental framework to profile the induced viral transcriptome. Subsequently, we employ an ONT-cappable-seq approach to map the full SSV1 transcriptome at base pair resolution, allowing a comprehensive characterization of its transcriptional architecture and regulatory characteristics.

## Results

### Mild UV exposure strongly increases SSV1 genome copy number

To identify conditions that promote active viral replication, transcription and transcriptional regulation while minimizing detrimental effects on host cell growth and survival, we investigated the response of SSV1-infected *S. solfataricus* S441 to UV exposure (**Figure 1** and **Table 1**). This was performed under conditions comparable to a previously proposed standardized protocol (19) (**Figure 1A**), and similar observations were made: increasing UV exposure markedly reduced cell growth, with exposure times of 45 seconds or higher and distances of 50 cm or lower strongly impairing cell growth and viability (**Figure 1B** and **Table 1**). The mildest condition tested, 30 seconds exposure at a 70-cm distance from the UV lamp, represented the least detrimental condition, with respect to growth (**Figure 1B**) and survival percentage (**Table 1**). For this condition, we performed quantification of virus presence, including both host-integrated and episomal viral DNA, using quantitative PCR (qPCR) on DNA extracted from UV-irradiated SSV1-infected *S. solfataricus* S441. This revealed an increase in viral DNA: at 12 hours after UV irradiation, the relative virus genome copy number showed a significant 100-fold increase (**Figure 1C**), which is within the same order of magnitude as previously observed (19).

**Figure 1.**
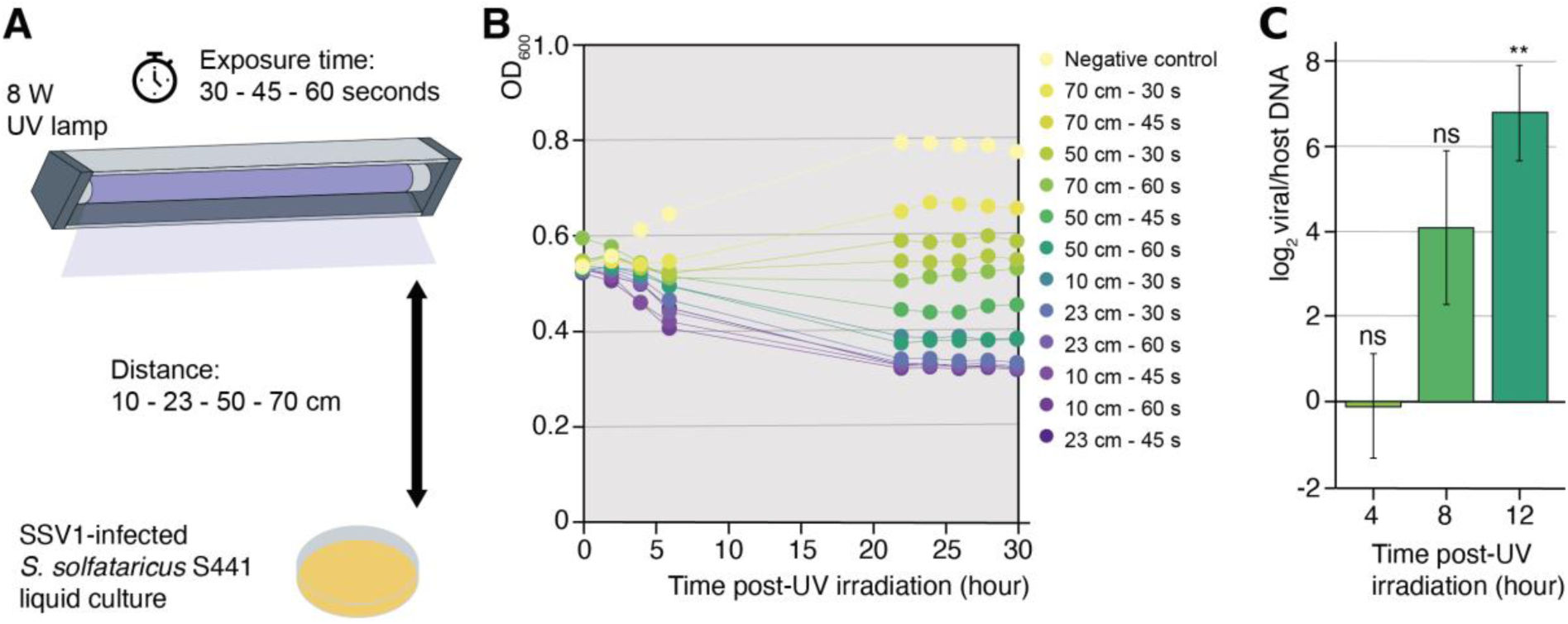
Effect of UV exposure on SSV1-infected *S. solfataricus* S441. (**A)** Schematic representation of the experimental set-up used for UV irradiation of liquid cultures. (**B**) Growth curves of liquid cultures post-UV irradiation. (**C**) Average fold change (Pfaffl R) in the ratio of viral/host DNA at three time points post-UV irradiation, relative to the 0-hour sample that was not subjected to UV treatment. Markers represent individual biological replicates. Columns represent mean Rs and error bars represent standard deviations (n=3). Log_2_(R) values were compared using a Student’s t-test. ** indicates p=0.0089, ns indicates no significant difference from 0 (R=1).

**Table 1.** Cell viability post-UV irradiation.

| Distance | Exposure time | CFU/mL | ± SD (n=3) | Survival percentage (%) |
| --- | --- | --- | --- | --- |
| 50 cm | 30 s | 7.7 × 10<sup>5</sup> | ± 1.5 × 10<sup>5</sup> | 10 |
| 50 cm | 45 s | 1.1 × 10<sup>5</sup> | ± 0.5 × 10<sup>5</sup> | 2 |
| 50 cm | 60 s | 0.4 × 10<sup>5</sup> | ± 0.5 × 10<sup>5</sup> | 1 |
| 70 cm | 30 s | 2.4 × 10<sup>6</sup> | ± 0.5 × 10<sup>6</sup> | 32 |
| 70 cm | 45 s | 9.3 × 10<sup>5</sup> | ± 1.2 × 10<sup>5</sup> | 12 |
| 70 cm | 60 s | 3.6 × 10<sup>5</sup> | ± 0.4 × 10<sup>5</sup> | 5 |
| Negative control |  | 7.5 × 10<sup>6</sup> | ± 1.1 × 10<sup>6</sup> | 100 |

### High-resolution transcription start site mapping reveals 5’-end diversity and promoter architecture

Next, we performed ONT-cappable-seq experiments with both UV-induced -employing the optimal UV-induction condition- and non-UV-induced (“carrier state”) cultures (**Figure 2**). Samples were pooled corresponding to early, middle and late time points post-UV induction (14). By capping the characteristic 5’-triphosphate groups of primary transcripts with desthiobiotin, enrichment can be performed using streptavidin beads. For each 5’-end peak, an enrichment ratio was calculated as reads per million (RPM) in the enriched sample divided by RPM in the control sample, in which no streptavidin-based enrichment was performed and all mRNA species were retained. Peaks with enrichment ratios above a set threshold were annotated as actual TSSs, distinguishing them from other possible 5’-ends originating from RNA processing (**Figure 2**; **Supplemental Table S1**).

**Figure 2.**
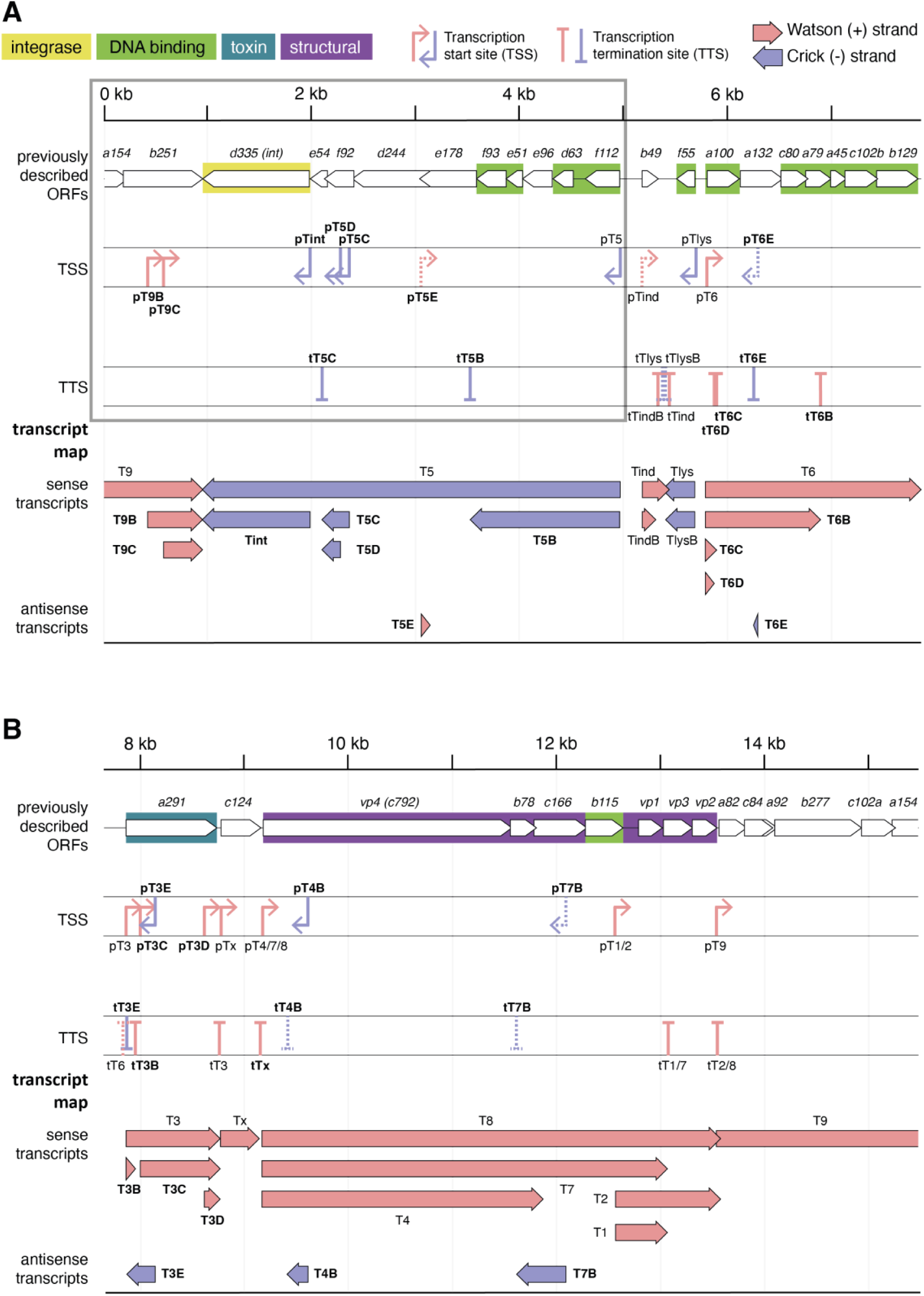
Linear representation of the full transcriptomic landscape of the SSV1 genome, including transcription start sites (TSSs) and transcription termination sites (TTSs) identified by ONT-cappable-seq data analysis. From top to bottom: SSV1 genome coordinates (13)(GenBank accession number NC_001338.1), previously described ORFs (8, 13), the full sets of TSSs and TTSs after ONT-cappable-seq, and the final transcript map with sense and antisense transcripts. ORFs are color-coded by the predicted function of their protein products (21). TSSs and TTSs newly identified in our study are labelled in bold, while TSSs and TTSs added after manual curation of the ONT-cappable-seq data are shown as dotted lines.

In total, 21 primary TSSs were identified across the SSV1 genome, displaying temporal regulation between carrier state and the UV-induction state at different times (**Figure 2**; **Supplemental Table S1**; **Supplemental Figures S1**-**S4**). Of these, 9 correspond to the described start sites of previously identified transcripts pT5, pTlys, pT6, pT3, pTx, pT4/7/8, pT1/2, pT9 and pTind (8, 11, 14), while 12 represent newly identified TSSs with condition-specific activity. Indeed, several of these newly identified TSSs were detected exclusively during early, middle and/or late stages following UV induction, whereas others were already active in the carrier state, indicating transcriptional reprogramming of the SSV1 genome during induction (**Supplemental Table S1**; **Supplemental Figures S1**-**S4**). For several transcript regions (T3, T5, T6 and T9), multiple TSSs were identified, with intragenic TSSs detected in addition to the primary TSS (**Figure 2**; **Supplemental Table S1**), suggesting transcriptomic plasticity and the generation of alternative 5’-end transcript isoforms.

In addition to sense TSSs driving transcription in the direction of annotated SSV1 genes, five antisense TSSs were detected through automated and/or manual analysis of the ONT-cappable-seq data: pT5E, pT6E, pT3E, pT4B and pT7B (**Figure 2**; **Supplemental Table S1**). Their detection in multiple conditions and in both enriched and control samples indicates that antisense transcription is a prevalent feature of the SSV1 transcriptome.

The final set of 21 SSV1 TSSs was compared to identify promoter signal elements and to assess whether promoter sequence features could explain condition-specific TSS detection (**Figure 3**). Putative BRE and TATA-box sequences were identified in promoter regions of most TSSs, except for pT5C, pT5E, pT3D and pT7B, where no canonical BRE/TATA could be identified. General trends across all promoter sequences include a clear TR initiator (Inr) dinucleotide around the TSS, an AT-rich PPE extending further upstream and a conserved TATA box around position −27 (**Figure 3A-B**). This TATA box is preceded by a weakly conserved BRE-like element, mainly reflected by two A/T bps at positions −35 and −34 (**Figure 3B-C**). These positions are shifted slightly upstream with the canonical Sulfolobales BRE consensus (15). Comparison of BRE and TATA-box signatures between promoters detected in all conditions (**Figure 3C**) and promoters detected only under specific UV-induction conditions (**Figure 3D-E**) did not reveal distinct sequence signatures associated with temporal promoter activity.

**Figure 3.**
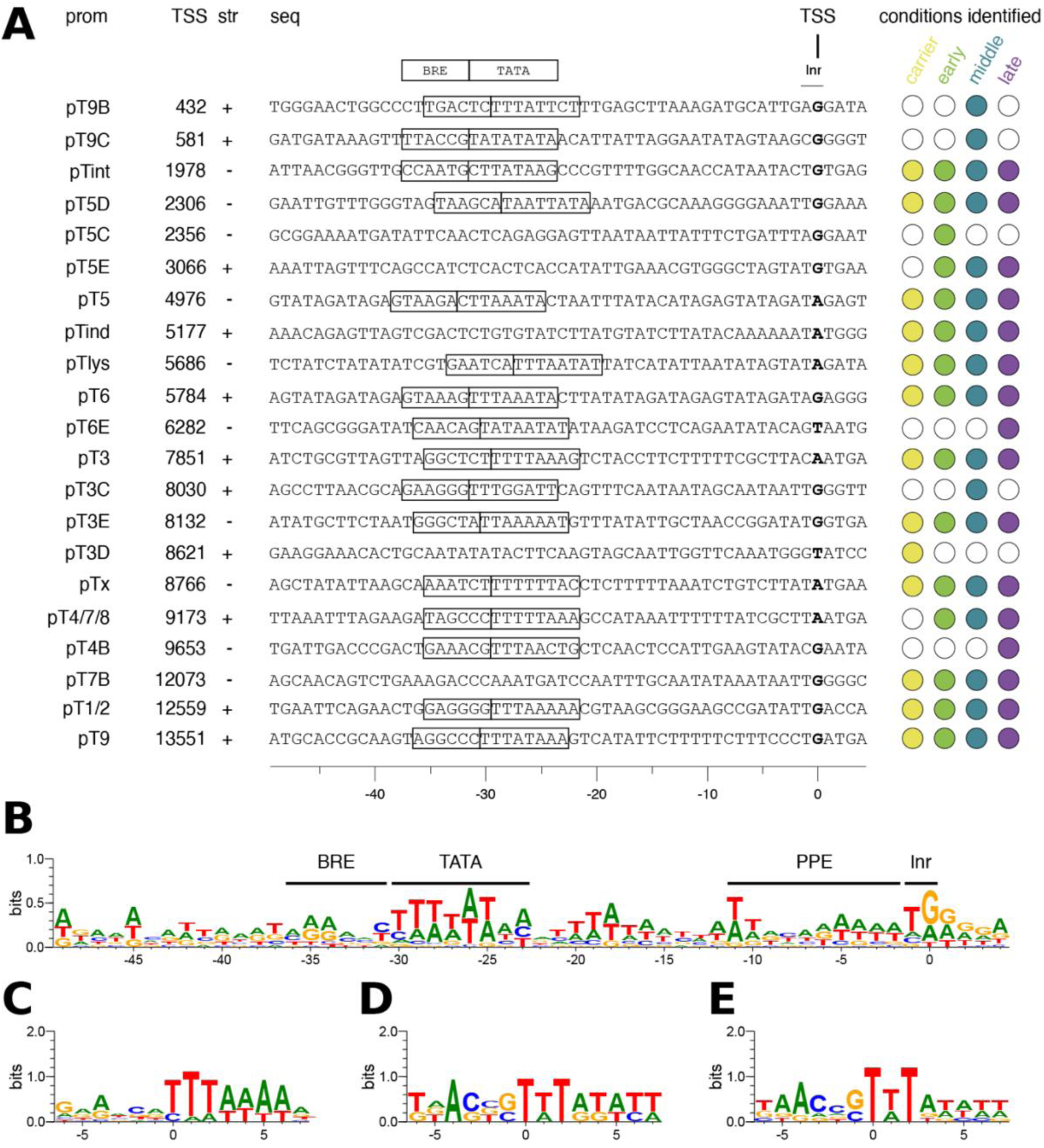
Sequence analysis on promoter regions of SSV1 Transcription Start Sites (TSSs). (**A**) Alignment of promoter regions preceding TSSs. For previously identified TSSs, BRE and TATA-box sequences are outlined as described in literature (8, 11, 13, 14). For newly identified TSSs, putative BRE and TATA sites have been annotated where possible, based on the previously determined BRE/TATA consensus sequence for Sulfolobales: RNWAAWTTTAWATR (15–17). Prom = promoter, str = strand, seq = sequence. (**B**) Sequence logo generated based on the alignment of promoter regions in A. (**C**) Sequence logo generated from alignment of BRE-TATA sequences of promoters identified in all conditions: pT5int, pT5D, pT5, pTlys, pT6, pT3, pT3E, pTx, pT1/2, pT9. pTind and pT7B could not be included because no BRE-TATA sequences were identified for these promoters. (**D**) Sequence logo generated from alignment of BRE-TATA sequences of promoters identified only at middle time points post-UV-induction: pT9B, pT9C, pT3C. (**E**) Sequence logo generated from alignment of BRE-TATA sequences of promoters identified post-UV induction: pT9B, pT9C, pT6E, pT3C, pT4/7/8, pT4B. pT5C, pT5E and pT3D could not be included because no BRE-TATA sequences were identified for these promoters. All sequence logos were generated using WebLogo v3.9.0 (https://weblogo.threeplusone.com/).

### Alternative transcription termination shapes 3’-end diversity

End-to-end nanopore sequencing also enabled identification of the original 3’-ends of primary transcripts, which were annotated as TTSs when the local drop in read coverage exceeded a defined threshold (**Figure 2**; **Supplemental Table S2**). In total, 19 TTSs were identified, of which 10 correspond to the previously described stop sites (8, 12, 13), while 9 have not been previously described. For several transcripts, including Tind and T6, multiple TTSs were associated with the same TSS, indicating alternative transcript 3’ ends and the formation of transcripts with different lengths (**Figure 2**). In contrast, mapped ONT-cappable-seq reads across all conditions tested revealed no clear stop sites for T4, T5 or T9. For T5 and T9, read coverage suggested extensive read-through from the T5 region into the downstream T9 region (**Supplemental Figures S1-S5**). For T4B, T6, T7B and Tlys, mapped ONT-cappable-seq reads clearly terminated in putative terminator regions (**Supplemental Figures S1-S4**). Therefore, despite being missing from the automated ONT-cappable-seq output, possibly due to RNA degradation, high terminator readthrough or low expression, they were included as TTSs (**Supplemental Table S2**).

Sequence analysis of the 19 identified TTSs showed that most stop sites are preceded by one or multiple oligo(dT)-stretches and a generally very AT-rich region, which is typical for *Sulfolobales* terminators (25, 26) (**Figure 4A-B**). No extended conserved terminator motif was apparent beyond this AT-rich context (**Figure 4B**). Typical oligo(dT) termination signals were also predicted in the respective terminator regions of T6 and Tlys. In contrast, no characteristic signals were identified in the sequences of the proposed terminator regions of T4, T5 or T9, consistent with the absence of clear stop sites for these transcripts in the ONT-cappable-seq data.

**Figure 4.**
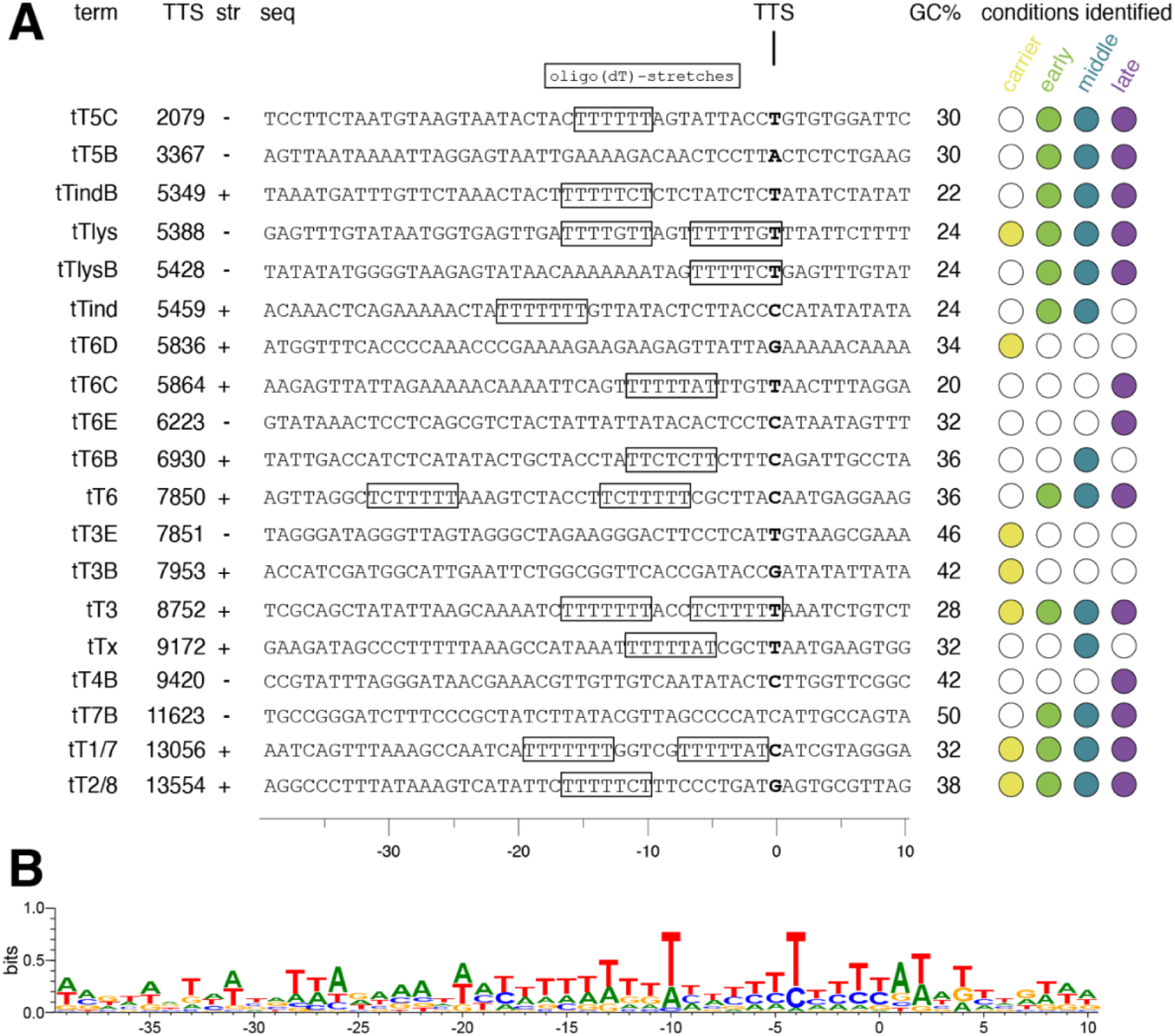
Sequence analysis on terminator regions of SSV1 Transcription Termination Sites (TTSs). (**A**) Alignment of terminator regions preceding TTSs. Oligo(dT)-stretches, which are typical transcription termination signals in Sulfolobales, have been annotated where present (18). Term = terminator, str = strand, seq = sequence, GC% = G/C percentage of terminator region. (**B**) Sequence logo generated based on the alignment of terminator regions in A, using WebLogo v3.9.0 (https://weblogo.threeplusone.com).

### Transcript unit reconstruction reveals nested, truncated and antisense transcripts

Combining information on TSSs and TTSs allowed subsequent delineation of transcription units (TUs) (**Figure 2**; **Supplemental Table S3**). Several newly resolved TUs were nested within previously described transcript regions, indicating that the SSV1 transcriptome is more modular than previously anticipated. For example, T9B and T9C were identified at the “middle” time point after UV induction (**Figure 5**, **Supplemental Figure S3**) and possibly encode ORFs of 510 and 294 nt, respectively, nested in-frame within *b251*, the terminal gene of T9 and a DnaA-like putative ATPase possibly involved in DNA packaging (27, 28).

**Figure 5.**
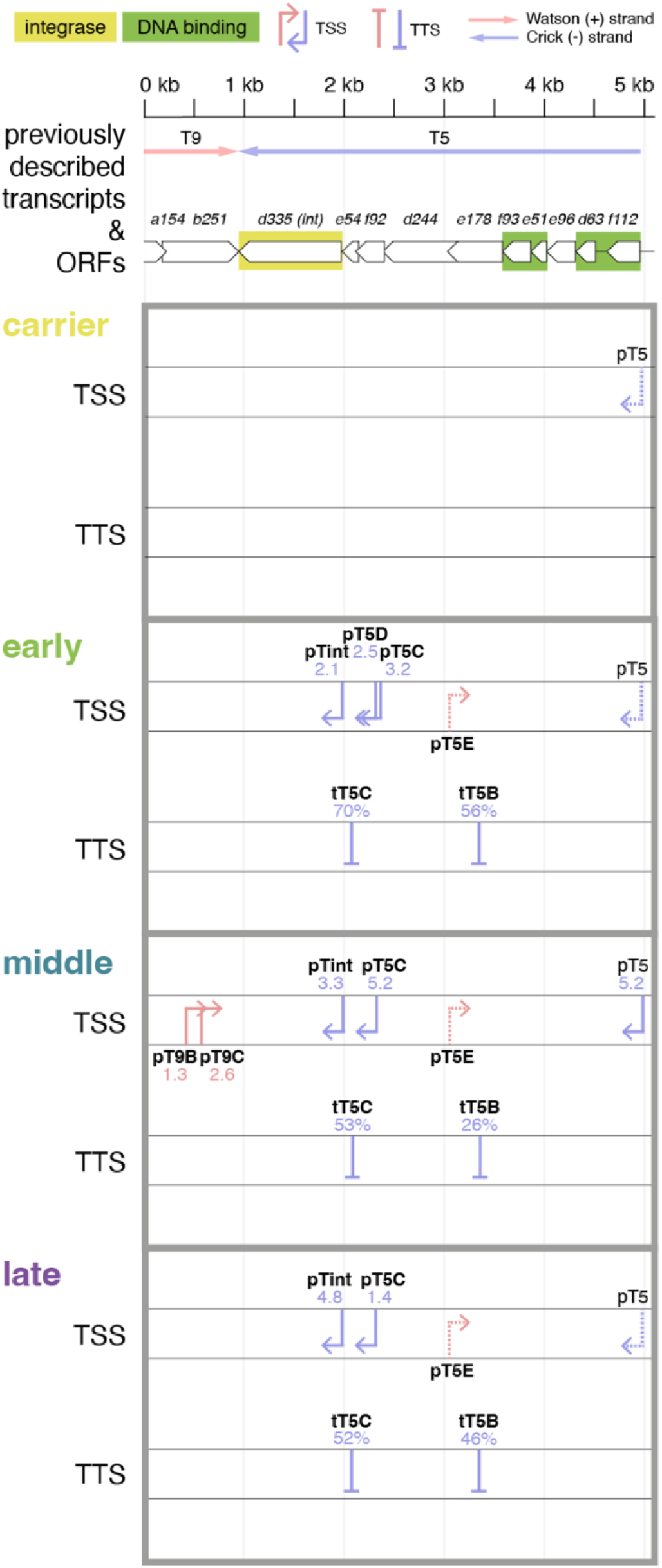
Overview of the T5 transcript region with TSSs and TTSs identified by ONT-cappable-seq for every condition tested: **carrier** state non-induced WT SSV1-infected *S. solfataricus* S441, **early** time points after UV-induction, **middle** time points after UV-induction, **late** time points after UV-induction. From top to bottom: SSV1 genome coordinates (13)(GenBank accession number NC_001338.1), previously described SSV1 transcripts (8, 11–14) and ORFs (8, 13), TSSs and TTSs identified in this region by ONT-cappable-seq data analysis, including the associated enrichment ratio for TSSs and the associated terminator efficiency for TTSs. ORFs are color coded by the predicted function of the protein products (21). TSSs and TTSs added after manual curation of the ONT-cappable-seq data are shown as dotted lines.

The T5 region provided a clear example of transcript diversification (**Figure 5**; **Supplemental Figure S5**; **Supplemental Table S3**). Within the previously identified T5 transcript of approximately 4016 nt, Tint was identified as a separate transcript encoding only the SSV1 integrase, the 3’-terminal gene on the larger T5 transcript, in both UV-induced and non-induced carrier-state conditions. Upstream of Tint, two even smaller nested transcripts were identified in the region containing the predicted *e54* and *f92* ORFs, which encode proteins of unknown function (29). However, little or no transcription spanning the complete *e54* and *f92* ORFs was observed, and together with the lack of clear promoter signatures for pT5C (**Figure 3A**), this suggests limited expression or possible misannotation of this SSV1-specific region. A shorter 3’-end-truncated variant was detected, T5B, which encodes the first five ORFs of T5, four of which are predicted DNA-binding proteins (21). This suggests that T5-derived transcripts may separate expression of putative regulatory genes from the remainder of the larger TU, including the integrase gene. Finally, transcription occurred antisense to ORFs *d244*-*e178* after UV induction, although no clear TTS was identified (**Figure 5; Supplemental Figure S5**).

The T3 region, containing a putative viral toxin gene *a291* (30), provided another example of transcript diversification with possible functional relevance (**Figure 2**). In addition to the main T3 transcript, T3C could encode an 84-amino acid truncated version of the A291 protein, whereas T3B represented a smaller carrier-state transcript, possibly resulting from premature termination (**Supplemental Figure S1**). Also in the carrier state, T3D was identified as the start of a relatively abundant ∼130-nt transcript at the 3’-end of T3, without an apparent ORF. Finally, the transcript T3E was detected antisense to the *a291* gene (**Figure 2**; **Figure 6**).

**Figure 6.**
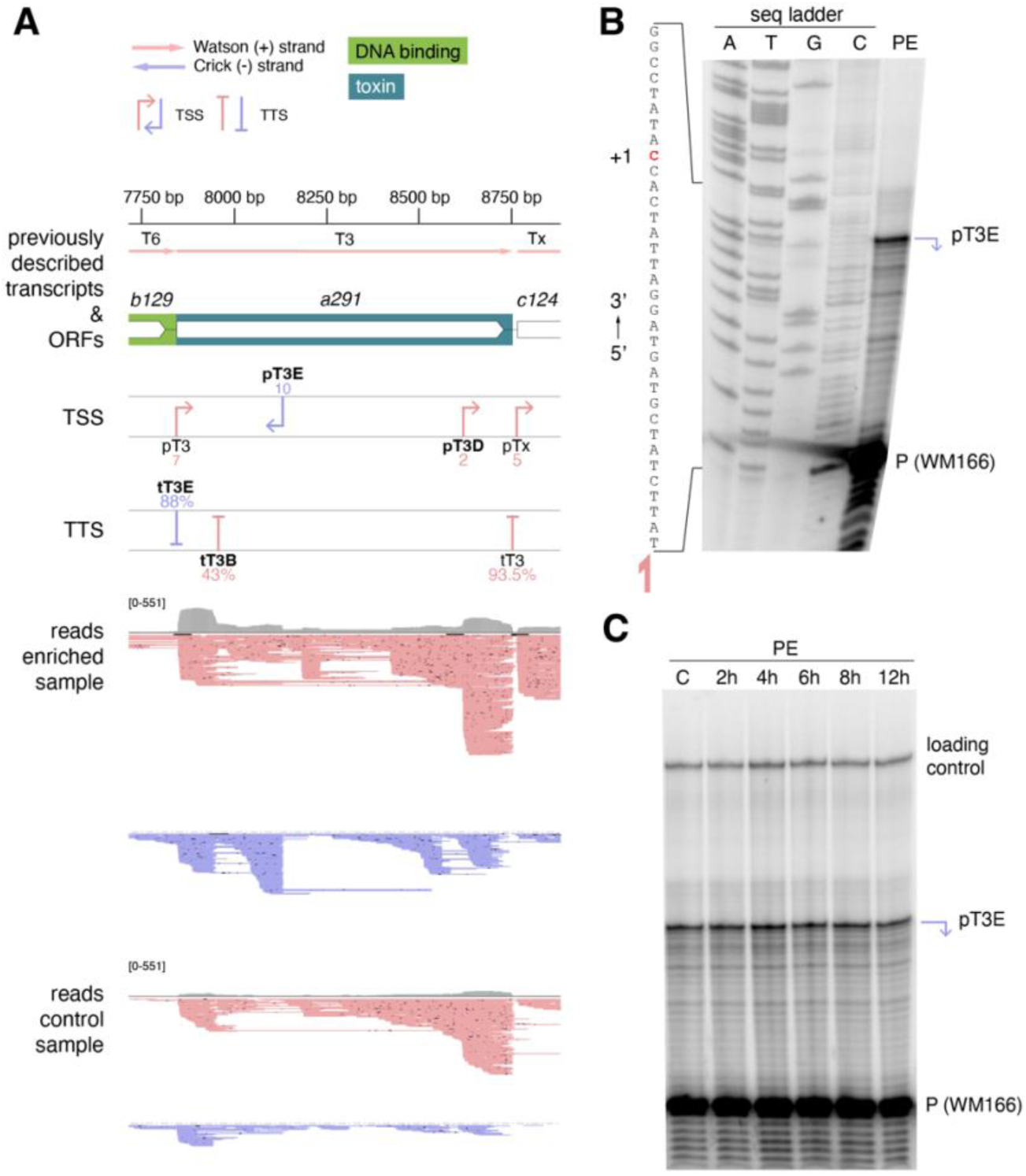
Detection of the T3E small RNA encoded antisense to the putative viral toxin *a291*. (**A**) ONT-cappable-seq results: close-up of the *a291* region for the “carrier state” sample, with T3E antisense reads between pT3E and tT3E. From top to bottom: SSV1 genome coordinates (13)(GenBank accession number NC_001338.1), previously described SSV1 transcripts (8, 11–14) and ORFs (8, 13), TSSs and TTSs identified in this region by ONT-cappable-seq data analysis, including the associated enrichment ratio for TSSs and the associated terminator efficiency for TTSs, and read coverage plotted in grey, with [0-551] being the y-axis scale. ORFs are color coded by the predicted function of the protein products (21). Black bars in read mapping indicate deletions/stretches of missing sequence in individual reads. (**B**) Experimental confirmation of pT3E transcription start site using primer extension on RNA extracted from WT SSV1-infected *S. solfataricus* S441 in non-induced “carrier state”. The first four wells (A, T, G, C) represent chain termination sequencing reactions. The full (sense) sequence is shown on the left. +1 = TSS. PE= primer extension product. P(WM166) = [γ-^32^P]ATP-labelled primer WM166, also represented with a pink arrow underneath the sequence on the left. (**C**) Primer extension using WM166 on RNA extracted from WT SSV1-infected *S. solfataricus* S441 at different time points post-UV-induction. Primer extension was performed on RNA-samples from 3 biological replicates (full picture in **Supplemental Figure S6**). A loading control (100 bp [γ-^32^P]ATP-labelled dsDNA) was added to each sample for normalization in densitometry calculations using ImageJ. PE = primer extension reactions. C= carrier state.

In addition to the 282-nt T3E, three more antisense transcripts were identified, ranging from 64 to 451 nt (**Supplemental Table S3**): T6E, overlapping antisense with respect to the *a132* ORF, T4B mapping antisense to *vp4* within the T4 region and T7B mapping antisense to *b78-c166* within the T7 region (**Supplemental Figures S2-S4**). While T7B was detected in all post-UV-induction samples, T4B appeared to be expressed only at late time points post-UV induction. The detection of these transcripts supports antisense transcription as a recurrent feature of the SSV1 transcriptome.

### A putative regulatory small RNA is encoded antisense to *a291*

Given its location antisense to the *a291* locus of T3, which encodes a putative viral toxin, T3E likely represents a novel small RNA (sRNA), potentially playing a regulatory role in the expression of T3 and thus A291 synthesis (**Figure 6A**). To experimentally validate this transcript, primer extension analysis was performed to confirm pT3E (**Figure 6B**). pT3E was detected in all conditions tested, including the non-induced carrier state and post-UV-induction condition (**Figure 6C**; **Supplemental Figures S6-S7**). Densitometric analysis showed that sRNA abundance did not differ significantly between conditions tested (**Supplemental Figure S8**). Moreover, RT-qPCR was performed to quantify relative expression ratios of antisense T3E to sense T3 and did not show significant up-or downregulation when comparing the post-UV-induction condition to non-induced carrier state (**Supplemental Figure S9**). Finally, secondary structure predictions suggested the formation of a stable RNA species (**Supplemental Figure S10**).

### Recombination occurs in SSV1 capsid protein genes

The ONT-cappable-seq analysis disclosed an unexpected feature in the SSV1 structural gene region in the genome (**Figure 7**): a substantial fraction of reads mapping to transcripts T2 and T8 contained a 286-nt gap in the structural protein-encoding genes *vp1* and *vp3* (**Figure 7A**). This gap corresponded exactly to the sequence between two 61 bp direct repeats that are present in the 3’-ends of both genes. From sequencing coverage data, around 20% of T2/8 transcripts were estimated to be missing this sequence in enriched samples and around 10% in non-enriched control samples, across all conditions tested (**Figure 7B-C**; **Supplemental Figure S11**). To investigate whether this deletion is located on the DNA- or RNA-level, PCRs were performed with cDNA generated from RNA samples collected before and after UV induction as well as with genomic DNA (gDNA) extracted in the same conditions (**Figure 7D-E; Supplemental Figure S12**). Both with cDNA and gDNA as a template, two differently sized amplicons were observed of 399 and 113 bp, corresponding to gDNA with and without the gap sequence, respectively. This indicates that the deletion was present on the genome level in a fraction of the viral DNA. VP1 and VP3, the SSV1 major and minor capsid proteins, respectively, are two homologous proteins (31), as shown by both structure and sequence alignments (**Figure 7F-G**).

**Figure 7.**
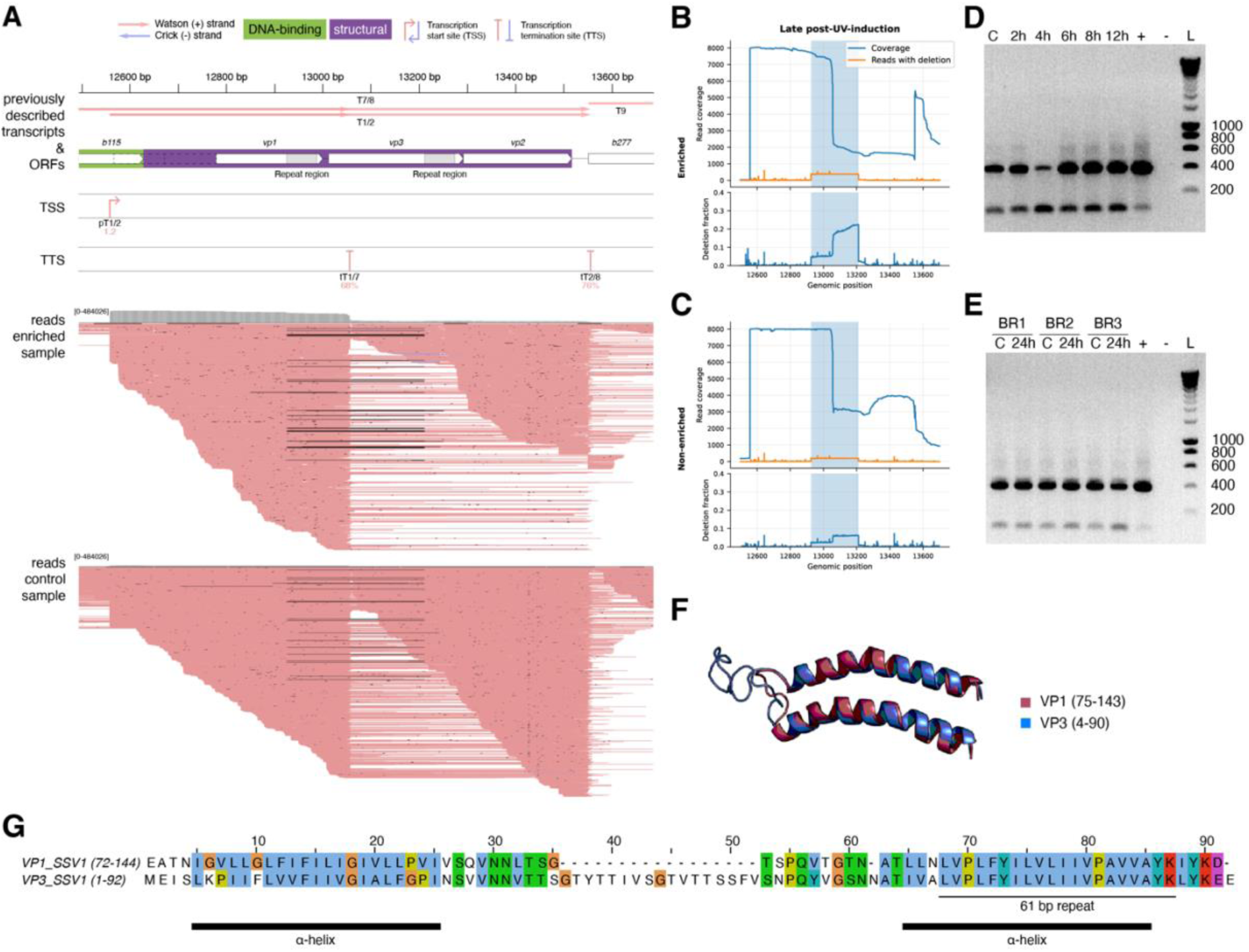
Recombination between direct repeats in SSV1 structural protein genes *vp1* and *vp3*. (**A**) ONT-cappable seq results: close-up of the SSV1 structural gene region for the “late” sample post-UV-induction. From top to bottom: SSV1 genome coordinates (13)(GenBank accession number NC_001338.1), previously described SSV1 transcripts (8, 11–14) and ORFs (8, 13), TSSs and TTSs identified in this region by ONT-cappable-seq data analysis, including the associated enrichment ratio for TSSs and the associated terminator efficiency for TTSs, and read coverage plotted in grey, with [0-484026] being the y-axis scale. ORFs are color coded by the predicted function of the protein products (21). The direct repeats in *vp1* and *vp3* are represented by grey boxes. For *vp1*, the white arrow matches the protein post-processing, with dotted lines indicating potential full-length versions of the gene from the 2 described possible start codons (31). Black bars in read mapping indicate deletions/stretches of missing sequence in individual reads. (**B** and **C**) Plots showing the read coverage (blue) and number of reads with a deletion (orange) (top graph), as well as the fraction of reads carrying a deletion (bottom graph) for every genomic position in the region 12500-13700, for both enriched (**B**) and non-enriched (**C**) samples of “late” timepoints post-UV-induction. The region 12926-13211 between the direct repeats is highlighted in blue. The same plots for all other conditions tested are shown in **Supplemental Figure S10**. (**D**) PCR on cDNA generated from RNA-samples extracted from WT SSV1-infected *S. solfataricus* S441 before and after UV-induction (only biological replicate 3 is shown, see **Supplemental Figure S11** for all biological replicates), and (**E**) PCR on DNA samples extracted from WT SSV1-infected *S. solfataricus* S441, either in carrier state (no UV-induction) or 24h post-UV-induction. For both PCRs expected products are 399 bp when the region between repeats is present, or 113 bp when the region between repeats is absent. BR = Biological Replicate. C = carrier state (no UV-induction). + = EAI_283 as stand-in SSV1 genome as positive control (29). - = H_2_O as negative control. (**F**) Structural alignment of VP1 (residues 75-143, in red) and VP3 (residues 4-90, in blue) proteins, performed using the Pairwise Structure Alignment tool on RCSB PDB (https://www.rcsb.org/alignment). (**G**) Protein sequence alignment of VP1 and VP3, performed using Clustal Omega (https://www.ebi.ac.uk/jdispatcher/msa/clustalo) and visualized using Jalview (https://www.jalview.org). In both structural and sequence alignments, the post-processing VP1 protein sequence was used. The residues corresponding to the 61 bp repeat region in both proteins are underlined, as well as the α-helix structural regions.

## Discussion

To understand transcriptional regulation on densely packed viral genomes, a high-resolution view of transcript boundaries is essential. ONT-cappable-seq has previously proven to be powerful for resolving TSSs, TTSs, associated promoter and terminator regions, operon structures and regulatory RNAs in bacteriophages (24, 32–37). Here, applying this approach to SSV1-infected *S. solfataricus* before and after UV induction revealed a more modular and dynamic transcriptional organization than suggested by the previously described set of 12 transcripts (8, 11–14).

The large “early” T5 transcript (14) provides a clear example of this modularity and how this might be linked to functional segregation. T5 is transcribed as three smaller, individual transcripts, with one (Tint) containing the integrase gene. Independent expression of the integrase has previously been observed for other *Fuselloviridae* (38, 39) and has been suggested for SSV1 (14, 29). Expression of Tint was more pronounced in a later stage post-UV-induction, which suggests that integrase expression is regulated independently from the full-length T5 transcript. This may be linked to a need for recombination later in the SSV1 life cycle, as the integrase promotes integrative and excisive recombination, relaxation of supercoiled DNA and recombination between two viral attP or two host attB sites (40–42). Additional T5-derived transcripts (T5C/T5D) further suggest that the larger T5 unit can be subdivided into functionally distinct modules, potentially separating expression of predicted regulatory genes from poorly characterized SSV1-specific ORFs (29, 43–46).

The transcript map did not only provide insight into promoter usage, but also revealed transcription termination as an additional regulatory layer. No clear transcription termination sites were identified for transcripts T4, T5 and T9, which is consistent with previous studies where only approximate 3’-ends were determined (12, 13). For T4, this may indicate that the mature RNA results from processing of the larger T7/T8 transcripts rather than from canonical transcription termination. For T5 and T9, we propose that instead of termination at *bona fide* terminator sequences, transcription could be terminating due to head-on collisions of convergently transcribing transcription elongation complexes. This mechanism of RNAP stalling and abortion of transcription elongation through collisions has been observed before in *E. coli,* yeast and bacteriophages and has been linked with transcription regulation (47, 48). Conversely, the identification of TTSs within 100 bp downstream of the T6 and T3 TSSs suggests that premature termination or abortive elongation may contribute to transcription control, as previously described for archaeal transcription regulation (48).

Transcription termination of the neighboring transcripts Tind and Tlys appears to be controlled by multiple consecutive terminators, a common and conserved strategy in Archaea, which can generate alternative 3’-UTRs, thereby creating opportunities for post-transcriptional regulation (25). The 3’-ends of both transcripts overlap and consequently, the central region seems to function as a bidirectional terminator with abundant oligo(dT)-stretches and a generally low GC-content. Moreover, the Tind-Tlys region is rich in palindromic and repeat sequences and has previously been hypothesized to contain the SSV1 origin of replication (14, 29). Together, these features suggest that transcription termination, RNA 3’-end formation and replication-associated sequence architecture may be functionally linked in this region.

Considerable antisense transcription was observed across the SSV1 genome, with five new antisense transcripts identified. It was previously reported that *S. solfataricus* encodes high numbers of non-coding RNAs, with *cis*-encoded antisense transcripts for at least 8% of all genes (16), making a similar trend in viruses infecting *S. solfataricus* plausible. The sense transcript T3, which encodes the putative viral toxin A291 (30), is expressed at high level (14), which was reflected in its high read coverage. Although no differential expression post-UV-induction could be shown for the antisense T3E transcript, it is tempting to speculate whether T3E could serve as an antitoxin-like RNA by regulating the expression of A291 from the T3 transcript. To our knowledge, no such Type I toxin-antitoxin system has been discovered in Archaea yet.

Finally, the ONT-cappable-seq analysis also revealed a genomic recombination event: a subpopulation of the viral DNA was missing a 286-bp stretch between two identical direct repeat sequences. This recombination at the *vp1* and *vp3* loci, was previously suggested by Iverson and colleagues, when observing deletions of *vp3* after transforming *S. solfataricus* with transposon insertion mutants in the *vp3* gene and *vp1-vp3* intergenic region (29). It can be postulated that the recombination mutant produces a chimeric version of the VP1 structural protein, in which four C-terminal amino acids of the major capsid protein (MCP) are replaced by five C-terminal VP3 residues, with concomitant loss of the VP3 minor capsid protein. Because the main difference between VP1 and VP3 is the loop between α-helices, this recombination event could result in alternative assembly of the spindle-shaped virion. In recent cryo-EM studies of other spindle-shaped viruses, helical strands of MCPs have been shown to make up the virion bodies (49–51). In SSV19, hydrophobic patches in neighboring VP1 α-helices interact, while the loop between helices makes up the outer surface of the capsid (51). SSV1 mutants lacking *vp3* show aberrant, elongated morphology and are less fit than wild-type SSV1 (29). This loss of fitness could explain why *vp1-vp3* recombination occurs only in a subpopulation of viruses, and may reflect altered presentation of the interhelical loop in the outer capsid surface. In conclusion, the base-resolution transcript map presented in this study shows that SSV1 gene expression is shaped by dynamic transcription initiation, termination and post-transcritional regulation, while genomic variation provides an additional route toward viral phenotypic diversification.

## Materials and methods

### Saccharolobus solfataricus growth conditions

*Saccharolobus solfataricus* S441 infected with wild-type (WT) SSV1 virus was used for all experiments (13, 52). Liquid cultures were grown in a New Brunswick Innova44 shaking incubator at 75°C and 160 rpm in filter sterilized 1X DT medium (pH 3.5) (53) and cell optical density was monitored at 600 nm (OD_600_). DT plates were prepared by mixing prewarmed, filter-sterilized 2X DT medium (pH 2.6), supplemented with 20 mM MgSO_4_ and 7 mM CaCl_2_, with an equal volume of freshly prepared, autoclaved 1.4% w/v GelRite® (Duchefa Biochemie) and immediately pouring the medium into petri plates (53). Cells were grown on solid medium by adding 4 mL of prewarmed soft layer gel, being DT medium + 02% w/v GelRite® (Duchefa Biochemie), to 1 mL of culture and pouring the mixture on a DT plate. Plates were incubated at 75°C for 7 days.

### UV irradiation of Saccharolobus solfataricus cells, DNA extraction and real time quantitative PCR

WT SSV1-infected S441 was cultivated in liquid DT medium to OD_600_ 0.5 (mid-exponential phase), after which 20 mL aliquots were transferred to 95 mm diameter Petri dishes for UV-irradiation in a dark room under red light. An 8 W 254 nm UV lamp (Cleaver Scientific) was placed at the desired distance (23, 50 or 70 cm) and the plates were exposed to UV light for the desired duration (30, 45 or 60 seconds) at room temperature (19). For studying post-UV-irradiation growth and viability, cultures were either incubated at 75°C and 160 rpm for growth monitoring up to 30 hours post-irradiation, or serial dilutions were plated on DT plates to assess colony-forming units (CFUs). Samples for DNA extraction were taken from the liquid cultures at 4, 8 and 12 hours post-irradiation, centrifuged (3000 *g*, 4°C, 15 minutes) and submitted to phenol-chloroform extraction and ethanol precipitation (54)(**Supplemental Note S1**). Virus/host DNA ratios in each sample were determined using quantitative PCR (**Supplemental Note S1**)(55).

### Total RNA extraction and library preparation for ONT-cappable-seq

Carrier-state WT SSV1-infected S441 was cultivated until OD_600_ 0.5, 9.5-mL samples were taken, combined with 1.9 mL stop mix solution (95% ethanol, 5% phenol, saturated with 0.1 M citrate, pH 4.5) and flash-frozen in liquid nitrogen.

An identical culture was cultivated until reaching OD_600_ 0.6 and subjected to UV induction, with the lamp set at 70 cm for 30 seconds. 7 mL samples were collected at time points 1, 2, 3, 4, 5, 6, 7, 8.5 and 12 hours post-irradiation, combined with 1.4 mL stop mix and flash frozen in liquid nitrogen.

RNA was isolated using hot phenol extraction and subsequent ethanol precipitation, followed by DNaseI treatment (ThermoFisher Scientific) (**Supplemental Note S2**). RNA purity and concentration were measured spectroscopically on SimpliNano (Biochrom US, Inc.) and fluorometrically on Qubit 4 using RNA HS assay kit (ThermoFisher Scientific). RNA integrity was evaluated on Agilent 2100 BioAnalyzer using RNA 6000 Pico kit.

Post-UV-induction RNA samples were combined at equal mass into 3 pools: 1, 2, 3, 4 and 5 hours post-induction samples were pooled into one “early” sample; 6 and 7 hours post-induction samples were pooled into one “middle” sample; 8.5 and 12 hours post-induction samples were pooled into one “late” sample. For each pool, 5 µg of RNA was supplemented with 1 ng of control spike-in RNA (1.8 kb), which had been *in vitro* transcribed from FLuc control template using HiScribe T7 high yield RNA synthesis kit (New England Biolabs), DNaseI-treated, ethanol-precipitated and spin-column purified (56). Library preparation was further carried out as described previously (24, 33, 56). Following PCR barcoding and amplification, equimolar amounts of all samples were pooled in a combined 100 fmol library. Nanopore adapters were ligated and sequencing was performed on a P2Solo sequencing device (Oxford Nanopore Technologies) using a PromethION flow cell (R10.4.1) (24, 33, 56).

### ONT-cappable-seq data analysis

Data analysis was conducted as described extensively in (56), using the pipeline available on GitHub (https://github.com/LoGT-KULeuven/ONT-cappable-seq; v2.0.0). Briefly, reads were processed using the pychopper and cutadapt tools, mapped to the *S. solfataricus* S441 and SSV1 reference genomes (13, 52) using minimap2, and low-quality mapped reads were removed using samclip. Local maxima of 5’- and 3’-ends were identified and clustered into representative peaks. Configuration parameters for annotation of transcript boundaries are shown in (**Supplemental Note S3**). Genomic alignments were also visually inspected using Integrated Genomics Viewer (IGV)(57) and both TSS and TTS positions were curated manually. Transcription units (TUs) were delineated where TSS and TTS were located adjacently on the same strand and at least one ONT-cappable-seq read spanned the entire length of the candidate TU. Promoter and terminator sets were aligned based on TSS/TTS and visually inspected to identify promoter/terminator elements.

### Primer extension analysis

WT SSV1-infected S441 was grown and UV-induced as described above. After 2, 4, 6, 8 and 12 hours, 20-mL samples were taken, centrifuged (8 minutes, 4000 *g*, 4°C) and pellets flash-frozen in liquid nitrogen. RNA extraction was performed (**Supplemental Note S2**) and RNA concentrations were measured spectrophotometrically using NanoDrop (ThermoFisher Scientific). Primer extension using primers WM166, WM167 and WM168 (**Supplemental Table S4**) was performed as described extensively in (Baes et al., 2022), and results were visualized on denaturing polyacrylamide gels (**Supplemental Note S4**).

### Reverse transcriptase quantitative PCR for gene expression analysis

The same post-UV-induction RNA-extraction samples from WT SSV1-infected S441 were reverse-transcribed to cDNA using the GoScript Reverse Transcriptase kit (Promega) and used as template in qPCR using primer pairs WM154/WM155 and WM171/WM172 (**Supplemental Table S4**)(**Supplemental Note S5**).

### PCR-based confirmation of genomic recombination

Genomic DNA and cDNA samples from WT SSV1-infected S441 post-UV-induction were used as templates in PCR using GoTaq G2 Green Master Mix (Promega) and primer pair WM158/WM159 (**Supplemental Table S4**). After PCR (2 minutes at 95°C, 35 cycles of 30 seconds at 95°C, 30 seconds at 52°C and 30 seconds at 72°C, and a final 5 minutes at 72°C), DNA was analyzed on agarose gel electrophoresis. EAI_283 vector (Iverson et al., 2017) as stand-in SSV1 genome was used as positive control template and nuclease-free H_2_O (Sigma-Aldrich) as negative control template.

## Acknowledgements

This work was supported by the Research Foundation – Flanders (FWO) through grant 1172725N to W.M., by the POSSIBL project, iBOF/21/092 to E.P. and R.L. and by the Vrije Universiteit Brussel (Strategic Research Program SRP91).

## Data, metadata and code availability

Raw and processed sequencing data generated in this study have been deposited in the Gene Expression Omnibus (GEO) at NCBI under accession number GSE335716. Output sequencing data files, growth measurement data, qPCR/RT-qPCR source data and figure source data are available at https://doi.org/10.5281/zenodo.20543159. The complete ONT-cappable-seq analysis pipeline is available at https://github.com/LoGT-KULeuven/ONT-cappable-seq, version 2.0.0.

