## Supplemental Materials for "High-resolution transcriptome analysis reveals a multilayered and dynamic transcriptional architecture in the archaeal virus SSV1"

Magnus *et al*.

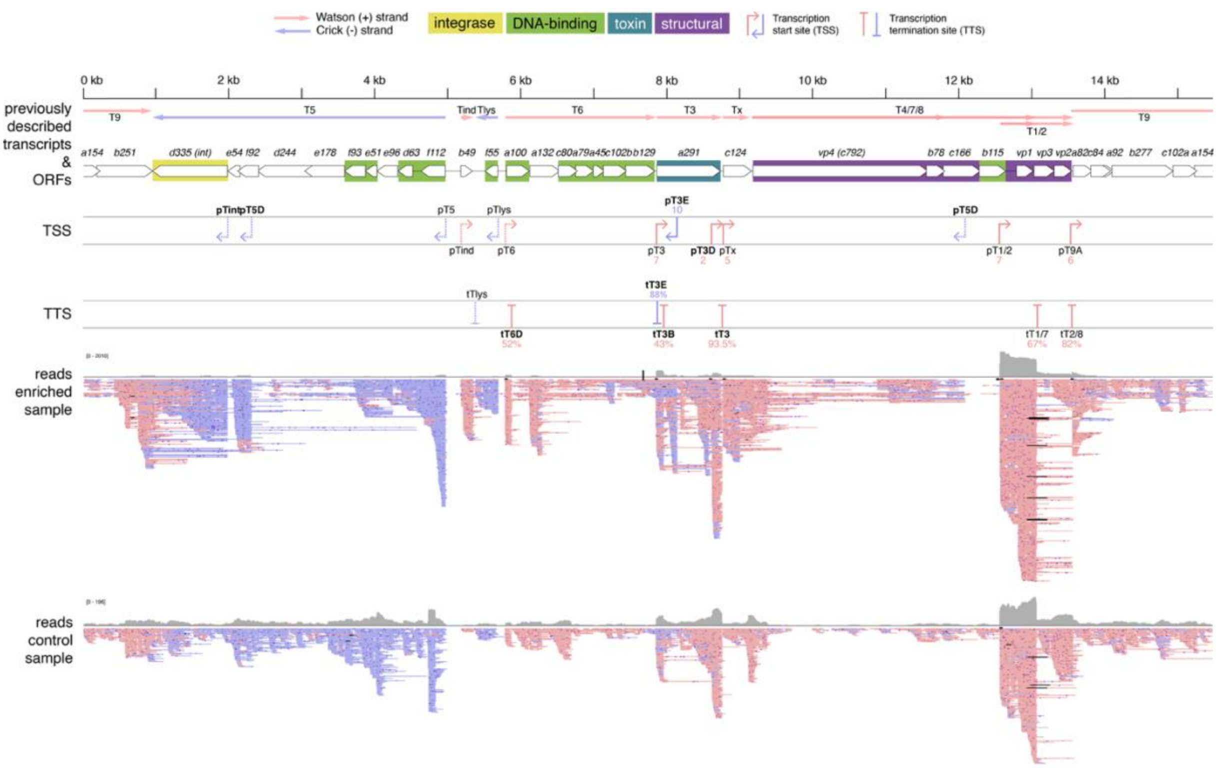


**Supplemental Figure S1.** Overview of mapped ONT-cappable-seq reads and transcript boundaries identified on SSV1 genome for carrier state samples. From top to bottom: SSV1 genome coordinates (1)(GenBank accession number NC_001338.1), previously described SSV1 transcripts (1–5) and ORFs (1, 5), TSSs and TTSs identified in this region by ONT-cappable-seq data analysis, and read coverage plotted in grey, with [0-2010/196] being the y-axis scale. ORFs are color coded by the predicted function of the protein products (6). TSSs and TTSs newly identified by ONT-cappable-seq are labelled in bold, while TSSs and TTSs added after manual curation of the ONT-cappable-seq data are shown as dotted lines. Black bars in read mapping indicate deletions/stretches of missing sequence in individual reads.

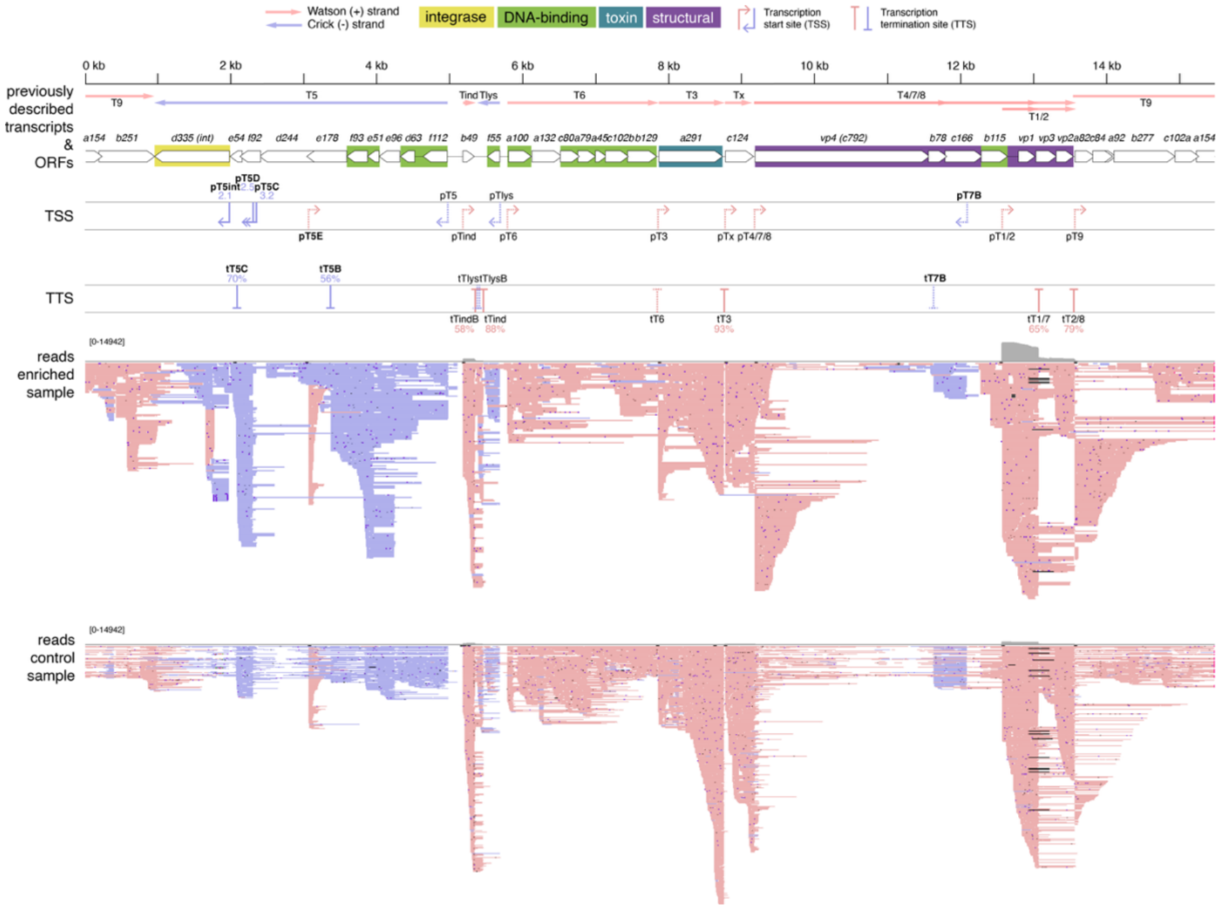


**Supplemental Figure S2.** Overview of mapped ONT-cappable-seq reads and transcript boundaries identified on SSV1 genome for early post-UV-induction samples. From top to bottom: SSV1 genome coordinates (1)(GenBank accession number NC_001338.1), previously described SSV1 transcripts (1–5) and ORFs (1, 5), TSSs and TTSs identified in this region by ONT-cappable-seq data analysis, and read coverage plotted in grey, with [0-14942] being the y-axis scale. ORFs are color coded by the predicted function of the protein products (6). TSSs and TTSs newly identified by ONT-cappable-seq are labelled in bold, while TSSs and TTSs added after manual curation of the ONT-cappable-seq data are shown as dotted lines. Black bars in read mapping indicate deletions/stretches of missing sequence in individual reads.

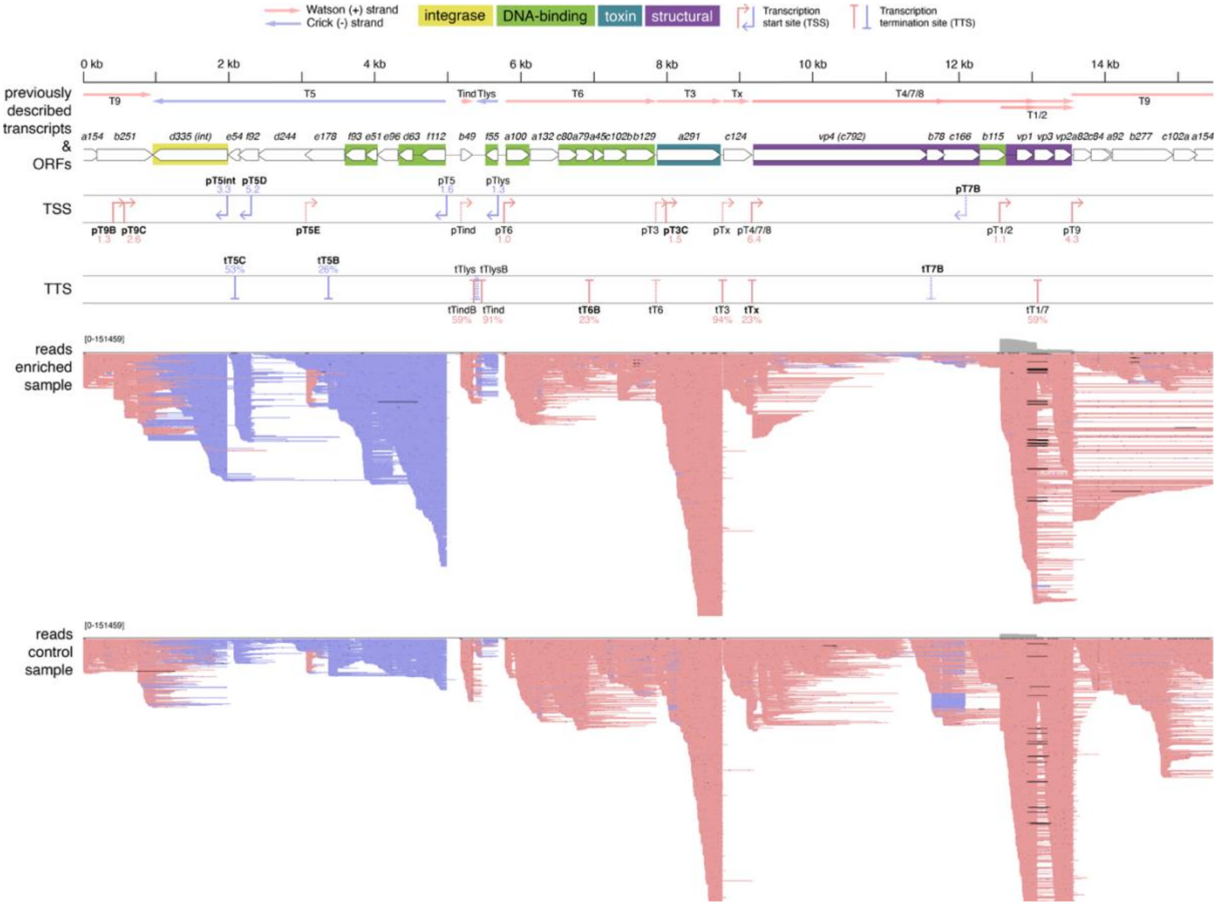


**Supplemental Figure S3.** Overview of mapped ONT-cappable-seq reads and transcript boundaries identified on SSV1 genome for <u>middle post-UV-induction</u> samples. From top to bottom: SSV1 genome coordinates (1)(GenBank accession number NC_001338.1), previously described SSV1 transcripts (1–5) and ORFs (1, 5), TSSs and TTSs identified in this region by ONT-cappable-seq data analysis, and read coverage plotted in grey, with [0-151459] being the y-axis scale. ORFs are color coded by the predicted function of the protein products (6). TSSs and TTSs newly identified by ONT-cappable-seq are labelled in bold, while TSSs and TTSs added after manual curation of the ONT-cappable-seq data are shown as dotted lines. Black bars in read mapping indicate deletions/stretches of missing sequence in individual reads.

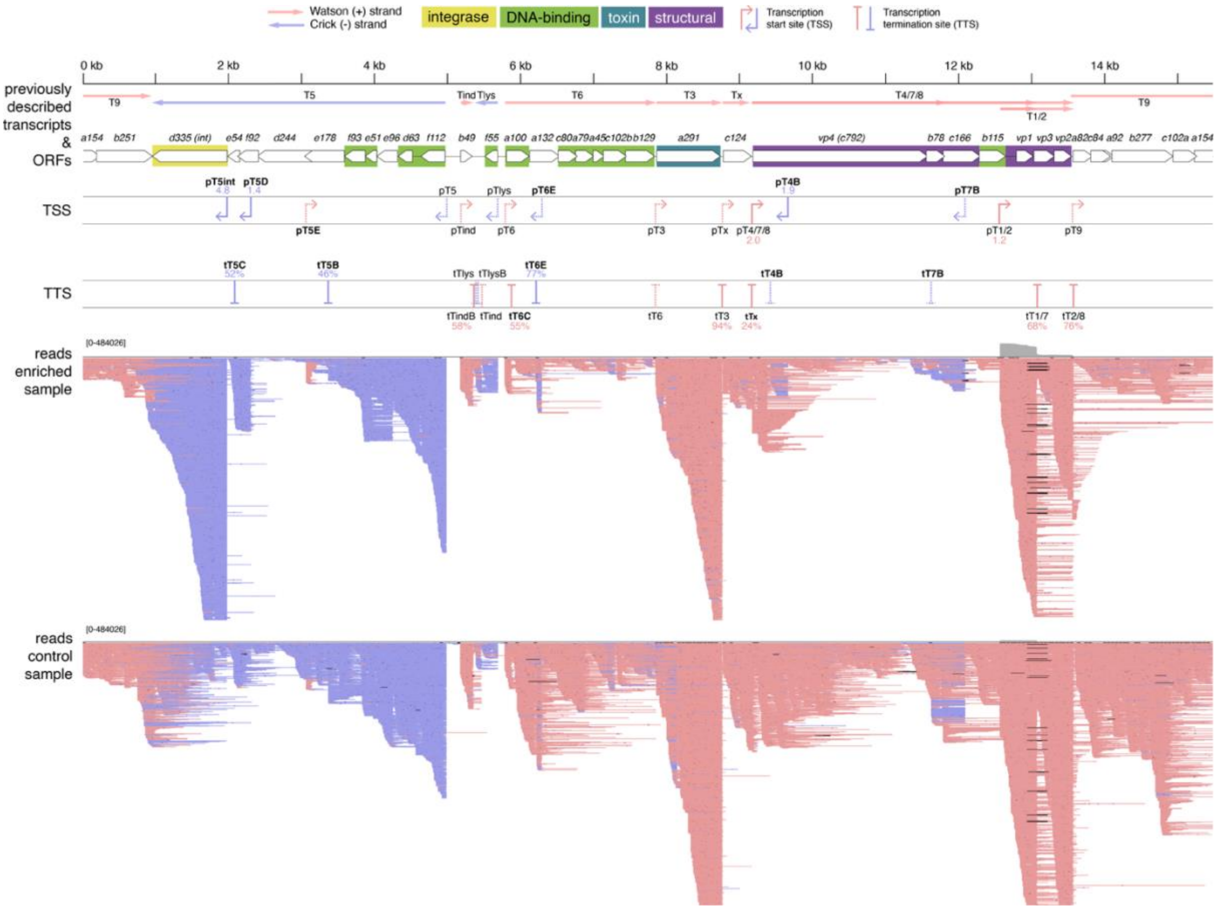


**Supplemental Figure S4.** Overview of mapped ONT-cappable-seq reads and transcript boundaries identified on SSV1 genome for late post-UV-induction samples. From top to bottom: SSV1 genome coordinates (1)(GenBank accession number NC_001338.1), previously described SSV1 transcripts (1–5) and ORFs (1, 5), TSSs and TTSs identified in this region by ONT-cappable-seq data analysis, and read coverage plotted in grey, with [0-484026] being the y-axis scale. ORFs are color coded by the predicted function of the protein products (6). TSSs and TTSs newly identified by ONT-cappable-seq are labelled in bold, while TSSs and TTSs added after manual curation of the ONT-cappable-seq data are shown as dotted lines. Black bars in read mapping indicate deletions/stretches of missing sequence in individual reads.

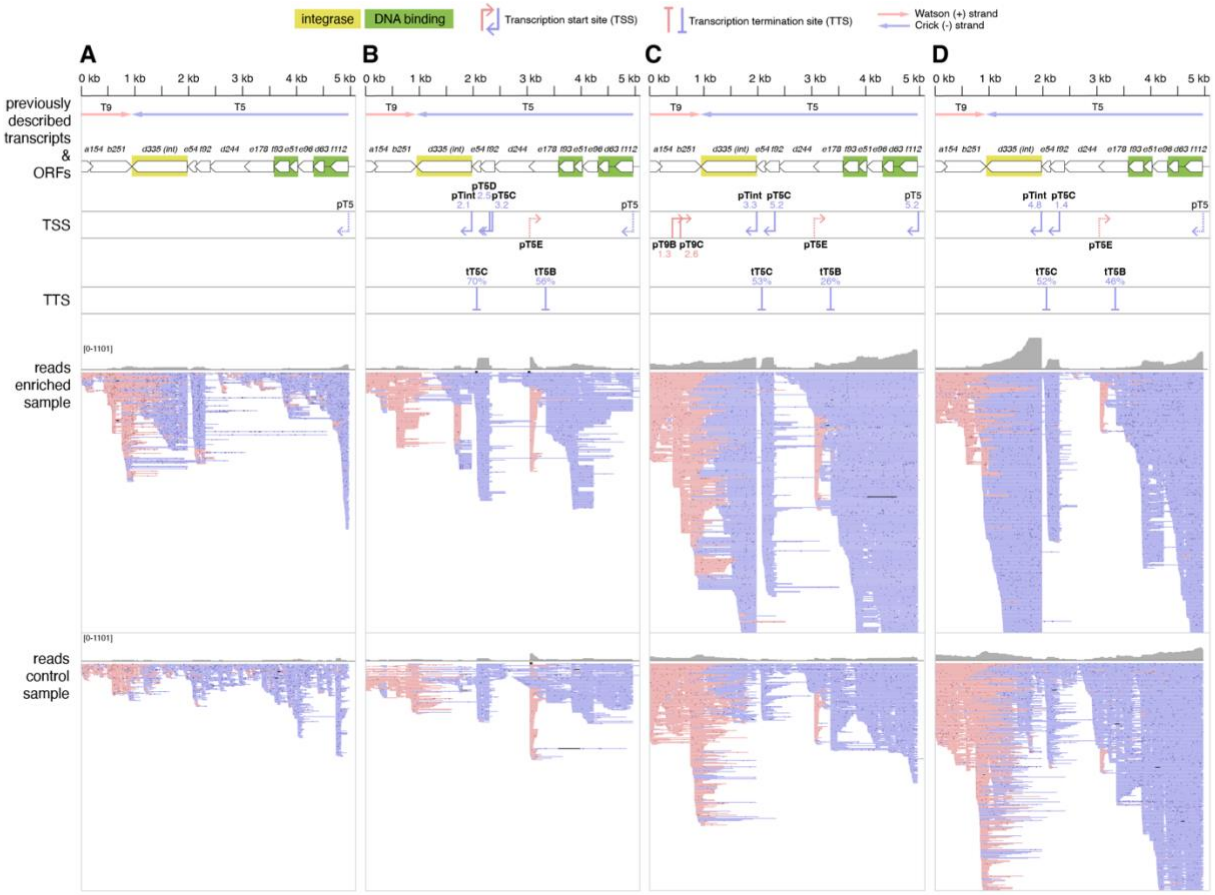


**Supplemental Figure S5.** Overview of the T5 transcript region with TSSs and TTSs identified by ONT-cappable-seq for every condition tested: (**A**) carrier state non-induced WT SSV1-infected S. solfataricus S441, (**B**) early time points after UV-induction, (**C**) middle time points after UV-induction, (**D**) late time points after UV-induction. From top to bottom: SSV1 genome coordinates (1)(GenBank accession number NC_001338.1), previously described SSV1 transcripts (1–5) and ORFs (1, 5), TSSs and TTSs identified in this region by ONT-cappable-seq data analysis, including the associated enrichment ratio for TSSs and the associated terminator efficiency for TTSs, and read coverage plotted in grey, with [0-1101] being the y-axis scale. ORFs are color coded by the predicted function of the protein products (6). TSSs and TTSs added after manual curation of the ONT-cappable-seq data are shown as dotted lines. Black bars in read mapping indicate deletions/stretches of missing sequence in individual reads. For “middle” and “late” conditions the y-axis was truncated at 1101 reads to allow better visualization of less abundantly covered regions.

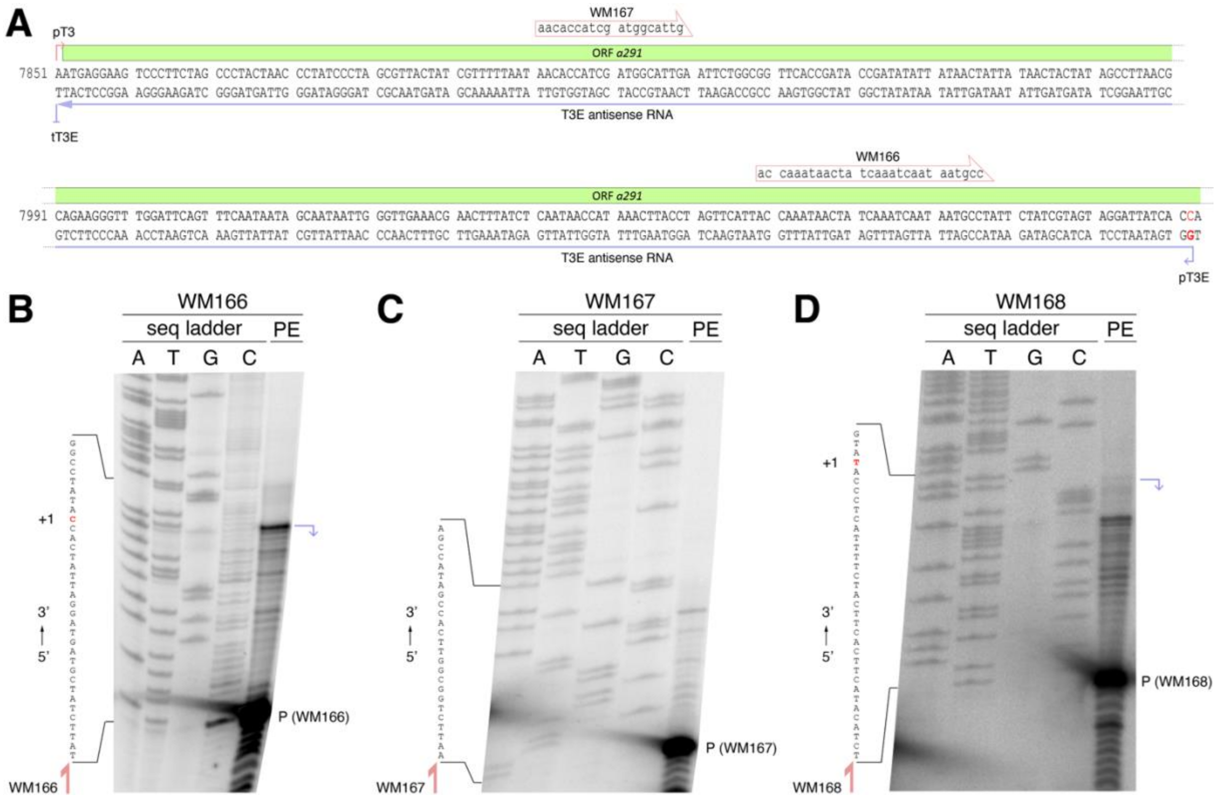


**Supplemental Figure S6.** Primer extension analysis using primers WM166, WM167 and WM168 on RNA-extraction samples from carrier state SSV1-infected *S. solfataricus* S441 culture samples. (**A**) Overview of the 7851-8133 region of the SSV1 genome sequence. TSSs pT3 and pT3E, TTS tT3E, ORF *a291*, T3E antisense transcript and the location of primers WM167 and WM168 are indicated. The GC basepair at position +1 representing TSS pT3E is highlighted in red. (**B**) Primer extension using WM166 for detecting TSS pT3E. (**C**) Primer extension using WM167 for detecting a possible intermediate TSS antisense from T3 between pT3E and tT3E. This was investigated given the peak of reads in this region in **Figure 6A**, but no intermediate TSS was detected. (**D**) Primer extension using WM168 for detecting the TSS for the *S. solfataricus porD* gene as mapped in (7). This was included to confirm the primer extension protocol used. The TSS as detected here differs slightly from (7), which could be due to the the use of *S. solfataricus* S441 used here as opposed to *S. solfataricus* P2 used in (7). For every gel, the 5’® 3’ sequence is shown on the left. The band corresponding to the [γ-$^{32}$P]ATP-labelled primers is indicated on the right. Purple arrows represent TSS pT3E in (**A**) and the TSS for the *porD* gene in (**B**) as identified in primer extension.

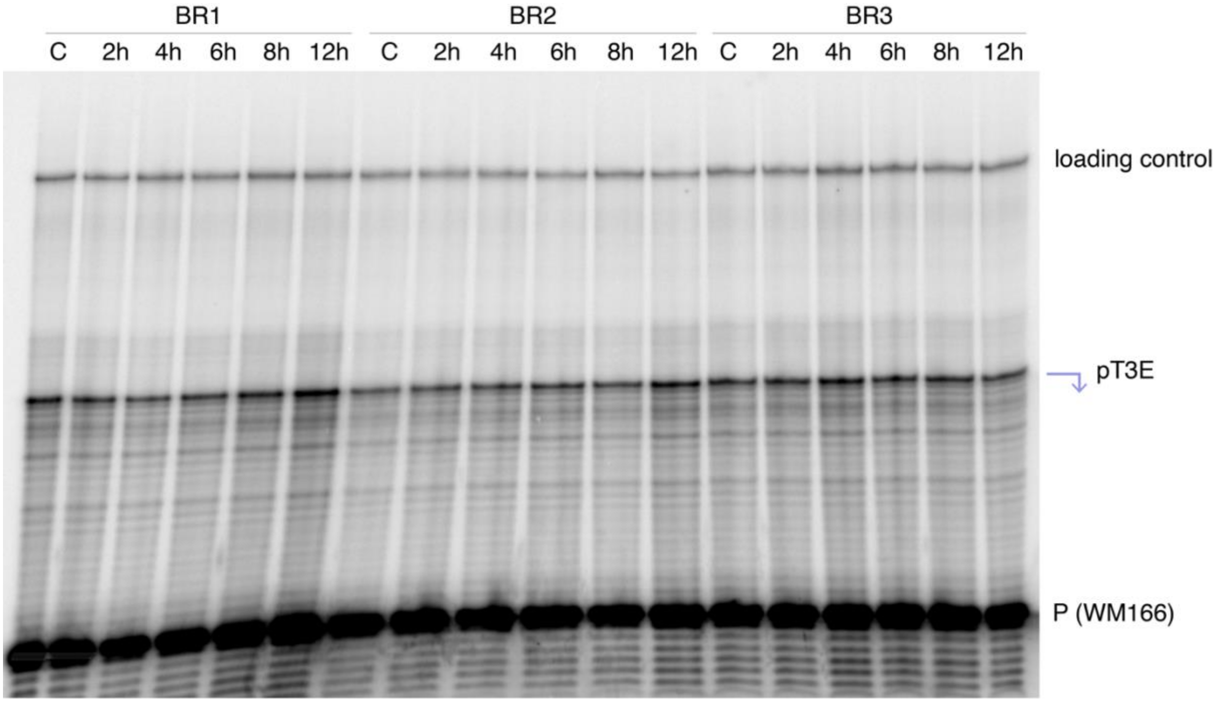


**Supplemental Figure S7.** Primer extension analysis mapping pT3E using primer WM166 on RNA-extraction samples from carrier state and post-UV-induction SSV1-infected *S. solfataricus* S441 culture samples, for 3 biological replicates. BR = biological control. C = carrier state. Loading control is a WM011-WM012 [γ-$^{32}$P]ATP -labelled dsDNA molecule, as described in Methods.

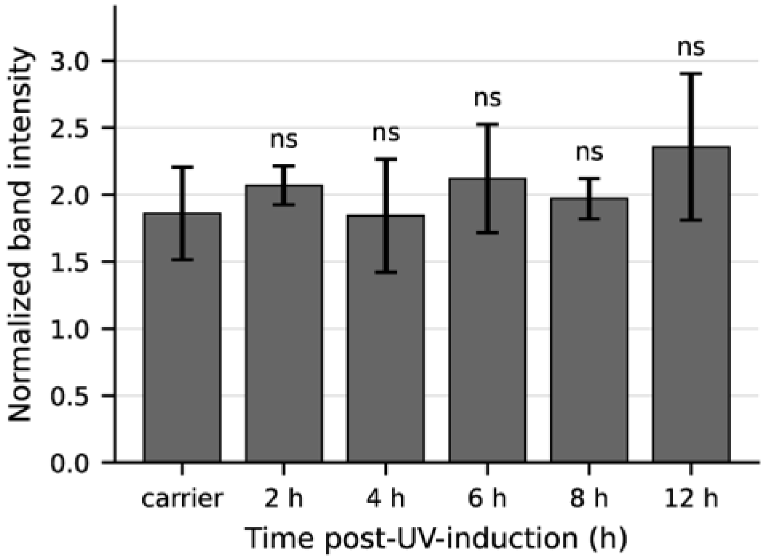


**Supplemental Figure S8.** Densitometric analysis on bands from primer extansion gel in Supplementary Figure S6. Band intensities were measured using ImageJ (https://imagej.net) and normalized for intensity of the loading control. Bars represent normalized band intensities averaged over 3 bilogical replicates. Error bars show standard deviations (n=3). Mean normalized band intensities for each time point were compared to carrier state using one-way ANOVA. No statistically significant differences (ns) among the six conditions were observed ($F(5,12) = 0.82$, $p = 0.56$).

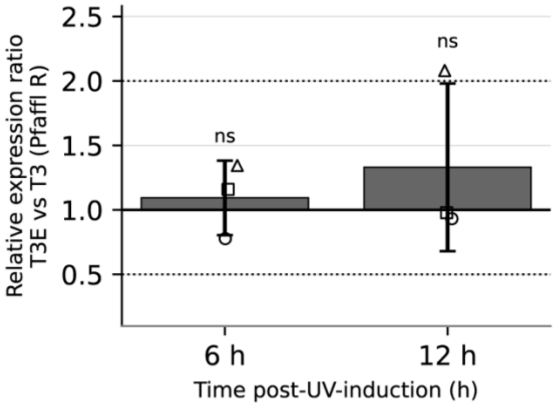


**Supplemental Figure S9.** RT-qPCR results comparing relative gene expression ratios of antisense T3E to sense T3 at different time points post-UV-induction. Average fold change (Pfaffl R) in gene expression ratio relative to 0 h (no UV-induction) are shown as bars, individual biological replicates are shown as markers and error bars represent standard deviations (n=3). Dotted lines indicate tresholds for 2.0 (upregulation) and 0.5 (downregulation). $Log_2$(R) values were compared using a Student's t-test. ns = no statistically significant difference.

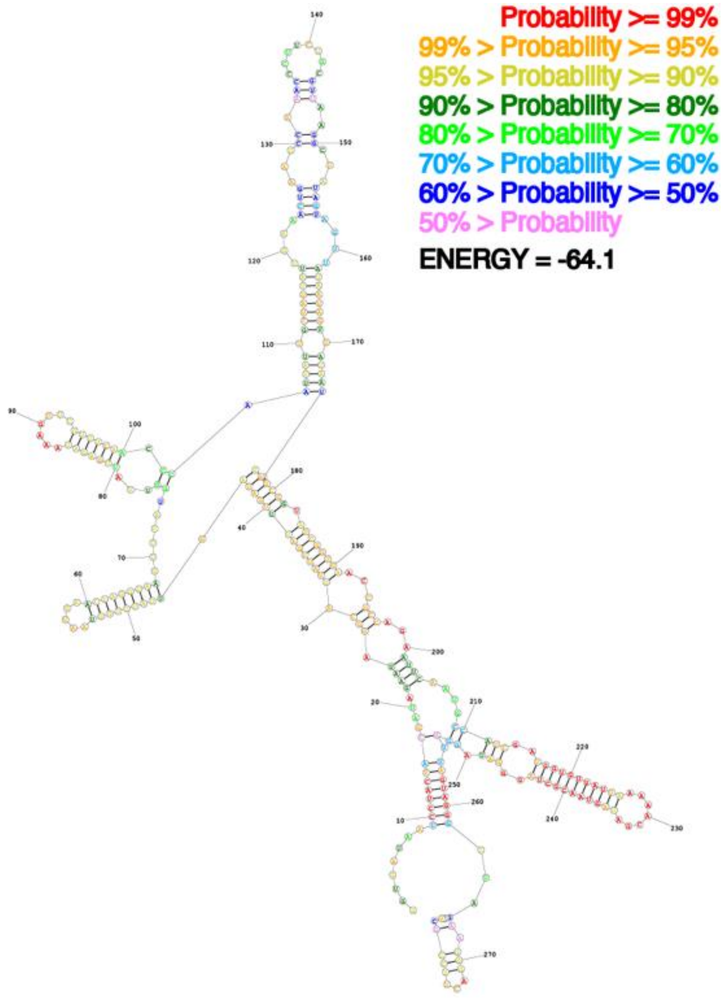


**Supplemental Figure S10.** Secondary structure prediction for the 282 bp-long pT3D-tT3C small RNA, using https://rna.urmc.rochester.edu/RNAstructure.html. The color legend representing base pairing probabilities and free energy estimation (in kcal/mol) is shown top right.

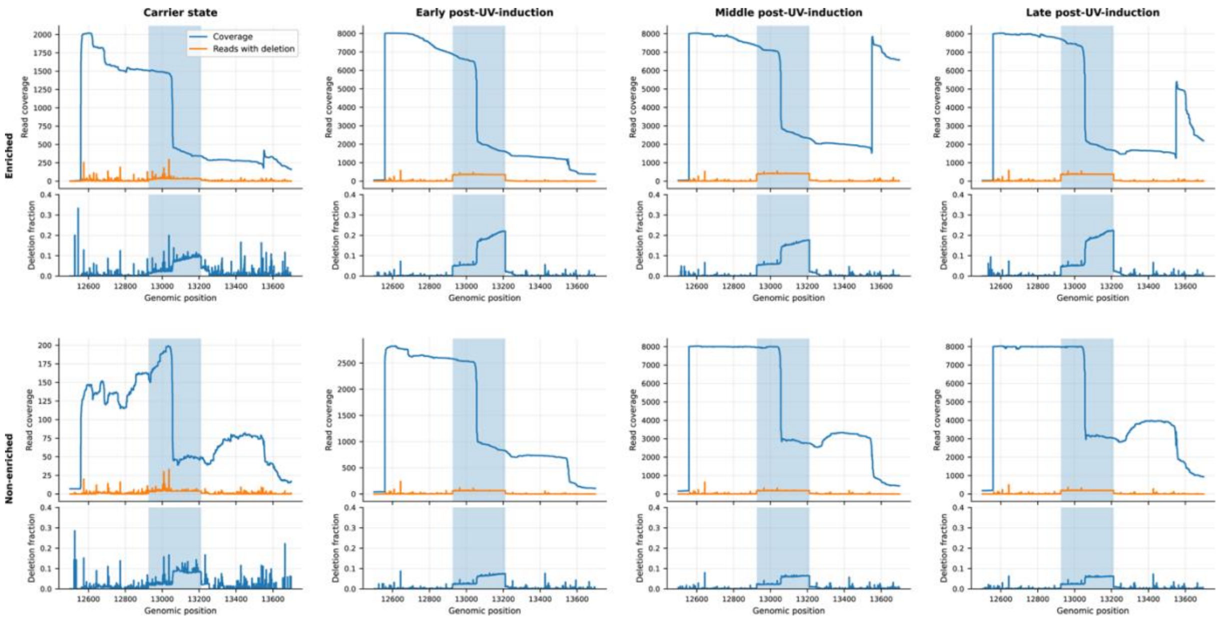


**Supplemental Figure S11.** Analysis of sequencing coverage and fraction of reads showing deletions for the SSV1 *vp1-vp3* structural gene region in all conditions tested (carrier state, early, middle and late post-UV-induction) and for both enriched and non-enriched samples. Plots showing the read coverage (blue) and number of reads with a deletion (orange) (top graph), as well as the fraction of reads carrying a deletion (bottom graph) for every genomic position in the region 12500-13700. The region 12926-13211 between *vp1-vp3* direct repeats is highlighted in blue.

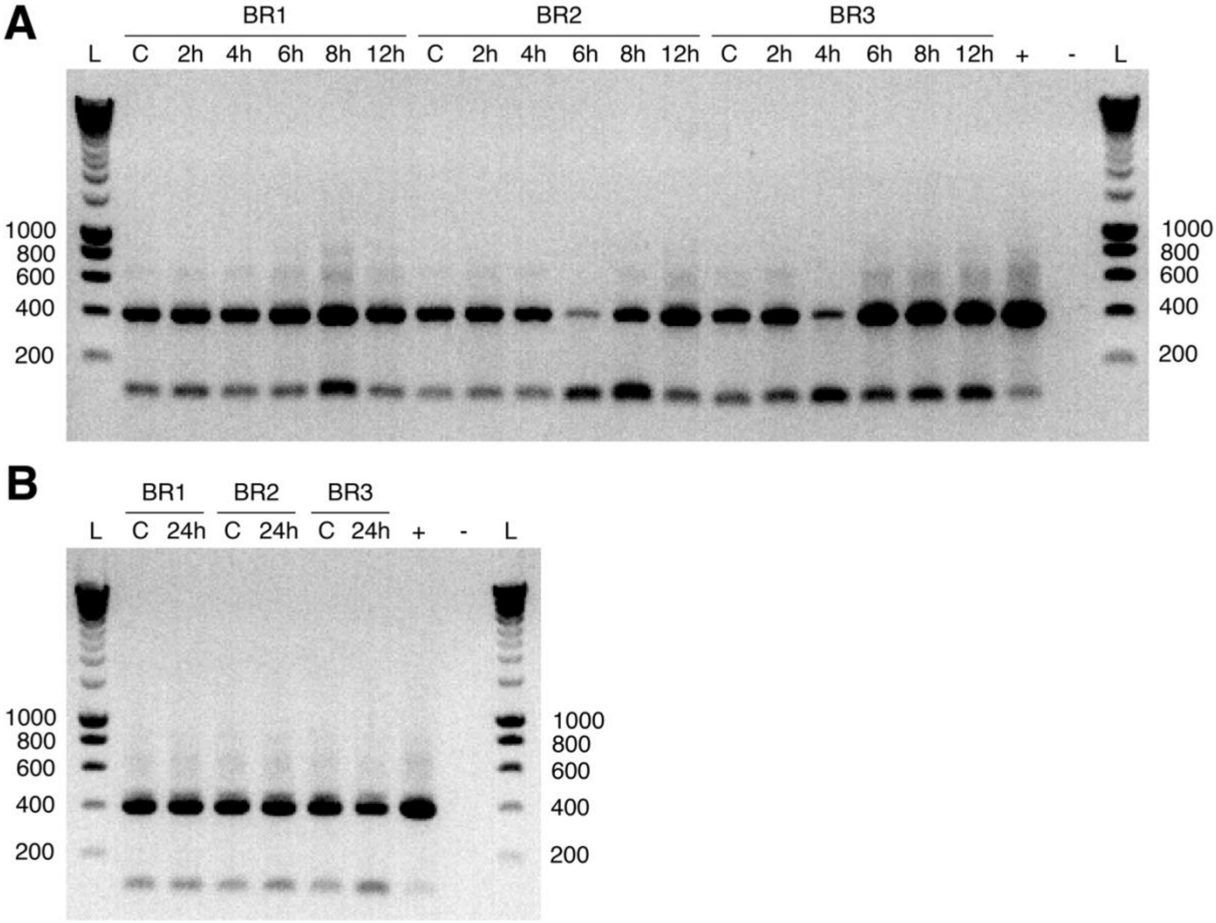


**Supplemental Figure S12.** (**A**) PCR on cDNA generated from RNA-samples extracted from WT SSV1-infected S. solfataricus S441 before and after UV-induction, and (**B**) PCR on DNA samples extracted from WT SSV1-infected *S. solfataricus* S441, either in carrier state (no UV-induction) or 24h post-UV-induction. For both PCRs expected products are 399 bp when the region between repeats is present, or 113 bp when the region between repeats is absent. BR = Biological Replicate. C = carrier state (no UV-induction). + = EAI_283 as stand-in SSV1 genome as positive control (8). - = $H_2O$ as negative control.

**Supplemental Note S1**. Detailed description of the DNA-extraction and real time quantitative PCR (qPCR) methods used for virus/host quantification post-UV-induction.

***DNA extraction from Saccharolobus solfataricus cells***

DNA extraction was performed using phenol-chloroform extraction and subsequent ethanol precipitation. Briefly, for each sample the cell pellet was resuspended in 200 µL 10 mM Tris-HCl, pH 8.5, to which an equal volume of phenol/chloroform/isoamyl (PCI) alcohol (25:14:1), pH 7.8-8.2 was added and the tube was vortexed vigorously for 1 minute. After 5 min centrifugation at 17000 *g*, the top aqueous layer was transferred to a new tube. 200 µL fresh 10 mM Tris-HCl, pH 8.5 was added to the PCI mix and the previous steps repeated, after which the top aqueous layer was combined with an equal volume of chloroform/isoamylalcohol 24:1, vortexed for 1 minute and centrifuged for 5 minutes at 17,000 *g*. The top aqueous layer was transferred to a new tube and combined with 0.75 M $NH_4OAc$, 20 µg glycogen and 2.5 volumes 100% ethanol. After overnight incubation at -20°C, the samples were centrifuged for 20 minutes at 4°C at 17000 *g*. The supernatant was removed and the pellet washed twice with 80% ethanol. After removing all ethanol, the DNA was resuspended in 10 mM Tris-HCl, pH 8.5.

***Real time quantitative PCR***

First, quantitative PCR (qPCR) oligonucleotide primers were tested for their amplification efficiency by running qPCR on purified EAI_283 plasmid DNA (8) as a stand-in SSV1 genome, employing SSV1-specific primers (vp2-fw, vp2-rv (9)), or purified *S. solfataricus* S441 genomic DNA, employing host-specific primers (WM026, WM027)(**Supplemental Table S4**). Template DNA was added in concentrations ranging from 0.1 pg/µL to 10.5 ng/µL in a reaction mix with 10 µL GoTaq qPCR MasterMix (Promega), 7.4 µL nuclease-free $H_2O$, 0.8 µL of each primer (10 µM) and 1 µL template. The qPCR program included 3 minutes at 95°C, followed by 40 cycles of 10 seconds at 95°C, 30 seconds at 55°C and the fluorescence measurement, after which a melting curve was measured by heating samples from 55°C to 95°C at a rate of 0.5°C/30 seconds. Primer efficiency was calculated by plotting threshold cycle (Ct) values against the log DNA concentration, performing linear regression and using the slope in the following equation: E = 10^-(1/slope). Off-target amplification was avoided by verifying that no extra peaks were present in the melting curve (-dRFU/dT vs T).

For qPCR analysis, DNA-extraction samples were used as a template in the same reaction mixture as described above for primer efficiency testing. The qPCR program setting was identical, except for the melting curve analysis. Virus/host DNA ratios were calculated using the Pfaffl mathematical model, in which the primer efficiencies (E) are used and the ΔCt (control vs sample conditions) of virus target is normalized with the ΔCt of host reference (Equation 1)(10).

$$R = \frac{{E_{virus}}^{\Delta C_{t,virus}(control-sample)}}{{E_{host}}^{\Delta C_{t},host(control-sample)}} \qquad (1)$$

Each biological replicate represents the mean of three technical replicates. Mean expression ratios and standard deviations were calculated from the three independent biological replicates. Statistical analysis was performed on $log_2$-transformed Pfaffl ratio’s using a Student’s t-test. A p-value below 0.05 was considered statistically significant.

**Supplemental Note S2**. Detailed description of RNA extraction using hot phenol and subsequent ethanol precipitation used for total RNA extraction for ONT-cappable-seq and primer extension.

Culture samples were thawed on ice, centrifuged for 20 minutes at 4°C and 4000 *g* and pellets were resuspended in TE buffer (10 mM Tris, 1 mM EDTA, pH 8) supplemented with 1% w/v sodium dodecyl sulphate (SDS). Samples were incubated at 64°C for 2 minutes to lyse the cells, after which 66 µL 3M NaOAc, pH 5.2 was added. 750 µL phenol was added and the samples were incubated at 64°C for 6 minutes, during which they were mixed multiple times by inversion. Samples were cooled on ice and centrifuged for 15 minutes at 17000 *g* and 4°C. The aqueous layer was removed and combined with 750 µL chloroform, mixed by inversion and centrifuged for 12 minutes at 17000 *g* and 15°C. The aqueous layer was again removed and combined with 1.4 mL of 30:1 EtOH:3M NaOAc, pH 6.5. The tubes were incubated at -20°C overnight and then centrifuged for 30 minutes at 17000 *g* and 4°C. The pellets were washed with 75% EtOH, and after removing all ethanol, the pellets were resuspended by adding 25 µL $H_2O$ and incubating in a thermomixer at 65°C for 5 minutes with gentle shaking. The resulting extraction samples were DNase-treated by combining 10 µL 4 µg/µL RNA with 5 µL DNaseI buffer, 4 µL (1 U/µL) DNase I (Thermo Fisher Scientific) and 0.5 µL RNaseOUT (ThermoFisher Scientific) followed by incubation at 37°C for 30 minutes. RNA was precipitated with ethanol as described above, followed by RNA Clean & Concentrator-5 spin-column purification kit (Zymo Research). Successful DNA removal was confirmed by DreamTaq PCR (ThermoFisher Scientific), using primers WM024 and WM025 (**Supplemental Table S4**).

**Supplemental Note S3**. Configuration parameters for identification of TSSs and TTSs in the ONT-cappable-seq data analysis pipeline.

```
termseq alpha: 0.001

cluster width:
  TSS: 15
  TTS: 30

minimum coverage:
  enriched:
    TSS: 25
    TTS: 25
  control:
    TSS: 2
    TTS: 5

peak alignment error: 2
TSS Threshold: 1
TTS threshold: 0.20

TSS sequence extraction:
  upstream: 50
  downstream: 1

TTS sequence extraction:
  upstream: 40
  downstream: 60
```

**Supplemental Note S4**. Detailed description of the primer extension protocol, based on (11).

Primers to be used for primer extension analysis (WM166, WM167, WM168) (**Supplemental Table S4**) were 5'-labelled with $^{32}P$ by adding 60 pmol primer to 25 μCi [γ-$^{32}P$]ATP and 10 units T4 Polynucleotide Kinase (PNK; ThermoFisher Scientific) in a 30 μL-reaction, incubating at 37°C for 40 minutes and quenching the reaction at 65°C for 10 minutes, yielding 2 pmol/μL $^{32}P$-labelled primer. DNA templates for generating sequencing ladders were amplified from the EAI_283 vector as stand-in SSV1 genome (8) using primers WM152 and WM113, and from genomic DNA extracted from WT SSV1-infected S441 (**Supplemental Note S1**) using primers WM169-WM170 (**Supplemental Table S4**). PCRs were performed using KAPA HiFi DNA polymerase (Roche) following manufacturer's instructions. PCR products were purified using phenol/chloroform extraction followed by ethanol precipitation (**Supplemental Note S1**). Sequencing ladders were prepared using the Thermo Sequenase Cycle Sequencing Kit (ThermoFisher Scientific). Briefly, 250 ng of the resulting purified DNA template was combined with 0.5 pmol labelled primer and 1 μL Thermo Sequenase DNA polymerase in a total volume of 8.75 μL sequencing master mix. For each dNTP, a tube was prepared containing 2 μL terminator mix (150 μM dNTP and 1.5 μM of the corresponding ddNTP) and 2 μL sequencing master mix. Single-stranded DNA synthesis was performed by incubating the tubes in a thermocycler for 55 cycles of 30 seconds at 94°C, 30 seconds at 55°C, 60 seconds at 72°C. The reaction was stopped by adding 2 μL formamide dye.

For primer extension, 10 μg RNA was combined with 4 pmol labelled primer in 22 μL total volume, incubated at 70°C for 5 minutes and chilled on ice for 5 minutes to ensure efficient hybridization. Reverse transcription was performed using GoScript Reverse Transcriptase (Promega) following manufacturer's instructions, with the only modification that the 1-hour extension step was performed at 55°C instead of 42°C, thereby preventing possible RNA secondary structures inhibiting the reverse transcriptase. cDNA was precipitated using ethanol (**Supplemental Note S1**) and resuspended in 6 μL formamide dye. Primer extension samples and corresponding sequencing ladders were run on a denaturing polyacrylamide gel, as described extensively in (11). After electrophoresis, read-out of the gels was performed using phosphorimaging.

Intensities of bands on polyacrylamide gels were measured using ImageJ (https://imagej.net). Band intensities for primer extensions were normalized to the intensity of a loading control: a 100 bp-long [γ-$^{32}P$]ATP -labelled dsDNA molecule obtained by labelling primer WM011, hybridizing it to its reverse complement WM012, and purifying the resulting dsDNA from gel as described in (12).

**Supplemental Note S5**. Detailed description of the reverse transcriptase real time quantitative PCR (RT-qPCR) method used for gene expression analysis.

Primer efficiency testing and qPCR on cDNA was performed as described above (**Supplemental Note S2**), using primer pair WM154-WM155 to target T3E, and primer pair WM171-WM172 to target T3 further downstream (**Supplemental Table S4**), outside the T3E region. Relative gene expression ratios were calculated using the Pfaffl mathematical model, in which the primer efficiencies (E) and the ΔCt ($C_t$ values in control *vs* sample conditions) for both targets were used (Equation 2)(10).

$$R = \frac{{E_{sRNA}}^{\Delta C_{t,1}(control-sample)}}{{E_{a291}}^{\Delta C_{t,2}(control-sample)}} \quad (2)$$

Each biological replicate represents the mean of two technical replicates. Mean expression ratios and standard deviations were calculated from the three independent biological replicates. Statistical analysis was performed on $\log_2$-transformed Pfaffl ratio's using a Student's t-test. A p-value below 0.05 was considered statistically significant.

**Supplemental Table S1**. List of the full set of SSV1 transcription start sites (TSSs) and associated promoter regions after ONT-cappable seq.

| TSS ID | TSS | +/- | Conditions identified | Annotation |
|---|---|---|---|---|
| pT9B | 432 | + | Middle post-UV-induction | Automatic |
| pT9C | 581 | + | Middle post-UV-induction | Automatic |
| pT5int | 1978 | - | Carrier state, early, middle and late post-UV-induction | Automatic + manual |
| pT5D | 2306 | - | Carrier state, early, middle and late post-UV-induction | Automatic + manual |
| pT5C | 2356 | - | Early post-UV-induction | Automatic |
| pT5E | 3066 | + | Early, middle and late post-UV-induction | Manual |
| pT5 | 4976 | - | Carrier state, early, middle and late post-UV-induction | Automatic + manual |
| pTind | 5177 | + | Carrier state, early, middle and late post-UV induction | Automatic + manual |
| pTlys | 5686 | - | Carrier state, early, middle and late post-UV induction | Automatic + manual |
| pT6 | 5784 | + | Carrier state, early, middle and late post-UV induction | Automatic + manual |
| pT6E | 6282 | - | Late post-UV-induction | Manual |
| pT3 | 7851 | + | Carrier state, early, middle and late post-UV induction | Automatic + manual |
| pT3C | 8030 | + | Middle post-UV induction | Automatic |
| pT3E | 8132 | - | Carrier state, early, middle and late post-UV induction | Automatic + identification in primer extension experiments |
| pT3D | 8621 | + | Carrier state | Automatic |
| pTx | 8766 | - | Carrier state, early, middle and late post-UV induction | Automatic + manual |
| pT4/7/8 | 9173 | + | Early, middle and late post-UV induction | Automatic + manual |
| pT4B | 9653 | - | Late post-UV induction | Automatic |
| pT7B | 12073 | - | Carrier state, early, middle and late post-UV induction | Manual |
| pT1/2 | 12559 | + | Carrier state, early, middle and late post-UV induction | Automatic + manual |
| pT9 | 13551 | + | Carrier state, early, middle and late post-UV induction | Automatic + manual |

**Supplemental Table S2**. List of the full set of SSV1 Transcription Termination Sites (TTSs) and associated terminator regions after ONT-cappable seq.

| TTS ID | TTS | +/- | Conditions identified | Annotation |
|---|---|---|---|---|
| tT5C | 2079 | - | Early, middle and late post-UV-induction | Automatic |
| tT5B | 3367 | - | Early, middle and late post-UV-induction | Automatic |
| tTindB | 5349 | + | Early, middle and late post-UV-induction | Automatic |
| tTlys | 5388 | - | Carrier state, early, middle and late post-UV-induction | Manual |
| tTlysB | 5428 | - | Early, middle and late post-UV-induction | Manual |
| tTind | 5459 | + | Early and middle post-UV-induction | Automatic |
| tT6D | 5836 | + | Carrier state | Automatic |
| tT6C | 5864 | + | Late post-UV-induction | Automatic |
| tT6E | 6223 | - | Late post-UV-induction | Automatic |
| tT6B | 6930 | + | Middle post-UV-induction | Automatic |
| tT6 | 7850 | + | Early, middle and late post-UV-induction | Manual |
| tT3E | 7851 | - | Carrier state | Automatic |
| tT3B | 7953 | + | Carrier state | Automatic |
| tT3 | 8752 | + | Carrier state, early, middle and late post-UV-induction | Automatic |
| tTx | 9172 | + | Middle and late post-UV-induction | Automatic |
| tT4B | 9420 | - | Late post-UV-induction | Manual |
| tT7B | 11623 | - | Early, middle and late post-UV-induction | Manual |
| tT1/7 | 13056 | + | Carrier state, early, middle and late post-UV-induction | Automatic |
| tT2/8 | 13554 | + | Carrier state, early, middle, and late post-UV-induction | Automatic |

**Supplemental Table S3**: List of SSV1 transcription units (TUs), delineated by ONT-cappable-seq TSSs and TTSs.

| TU | TSS | TTS | Length (nt) | +/- | ORFs encoded | Notes |
|---|---|---|---|---|---|---|
| T9 | pT9 | | ~2861 | + | *a82*, *c84*, *a92*, *b277*, *c102a*, *a154*, *b251* | No TTS identified |
| T9B | pT9B | | ~515 | + | *b251* (C-terminal truncated version) | No TTS identified |
| T9C | pT9C | | ~366 | + | *b251* (C-terminal truncated version) | No TTS identified |
| T5 | pT5 | | ~4016 | - | *f112*, *d63*, *e96*, *e51*, *f93*, *e178*, *d244*, *f92*, *e54*, *d335* (*int*) | No TTS identified |
| T5B | pT5 | tT5B | 1610 | - | *f112*, *d63*, *e96*, *e51*, *f93* | |
| T5C | pT5C | tT5C | 278 | - | *f92* (C-terminal truncated version) | |
| T5D | pT5D | tT5C | 228 | - | *f92* (C-terminal truncated version) | |
| Tint | pTint | | ~1018 | - | *d335* (*int*) | No TTS identified |
| T5E | pT5E | | variable | + | | Antisense, no TTS identified |
| Tind | pTind | tTind | 283 | + | *b49* | |
| TindB | pTind | tTindB | 173 | + | *b49* | |
| Tlys | pTlys | tTlys | 299 | - | *f55* | |
| TlysB | pTlys | tTlysB | 259 | - | *f55* | |
| T6 | pT6 | tT6 | 2067 | + | *a100, a132, c80, a79, a45, c102b, b129* | |
| T6B | pT6 | tT6B | 1147 | + | *a100, a132, c80* | |
| T6C | pT6 | tT6C | 81 | + | | |
| T6D | pT6 | tT6D | 53 | + | | |
| T6E | pT6E | tT6E | 64 | - | | antisense |
| T3 | pT3 | tT3 | 902 | + | *a291* | |
| T3B | pT3 | tT3B | 103 | + | | |
| T3C | pT3C | tT3 | 723 | + | | |
| T3D | pT3D | tT3 | 132 | + | | |
| T3E | pT3E | tT3E | 282 | - | | antisense |
| Tx | pTx | tTx | 407 | + | *c124* | |
| T8 | pT4/7/8 | tT2/8 | 4382 | + | *vp4 (c792), b78, c166, b115, vp1, vp3, vp2* | |
| T7 | pT4/7/8 | tT1/7 | 3884 | + | *vp4 (c792), b78, c166, b115, vp1* | |
| T4 | pT4/7/8 | | 2648 | + | *vp4 (c792), b78* | No TTS identified |
| T4B | pT4B | tT4B | 234 | - | | antisense |

| | | | | | | |
|---|---|---|---|---|---|---|
| T7B | pT7B | tT7B | 451 | - | | antisense |
| T2 | pT1/2 | tT2/8 | 996 | + | *vp1, vp3, vp2* | |
| T1 | pT1/2 | tT1/7 | 498 | + | *vp1* | |

**Supplemental Table S4**. Overview of primers used in this study.

| **Primer name** | **Sequence (5’ ® 3’)** | **F/R** | **Use** |
|---|---|---|---|
| Vp2-fw<br>Vp2-rv | GGAGGGTACATCGCTACCTTATGA<br>CAGTAGGGCTGACAGTAAACTACG | F<br>R | Virus-specific primers (targeting *vp2*) for quantification of relative virus copy numbers (9) |
| WM026<br>WM027 | TCTTCTCTCTTCGCTCCAGT<br>GGCAAGCCTAAAATCCAAATCC | F<br>R | Host-specific primers (targeting *S. solfataricus tbp*) for quantification of relative virus copy numbers |
| WM024<br>WM025 | ACAAAAGGAGGGGCGTTAGA<br>GGGATAGGGTTAGTAGGGCT | F<br>R | Verify absence of SSV1 DNA (targeting *b129*) in RNA-extraction samples for ONT-cappable-seq |
| WM166 | ACCAAATAACTATCAAATCAATAAT<br>GCC | F | Primer extension targeting TSS pT3E |
| WM167 | AACACCATCGATGGCATTG | F | Primer extension targeting potential alternative TSS for sRNA downstream from pT5C. |
| WM168 | CTGGAAGATACCTCTATACACTATC | F | Primer extension targeting *S; solfataricus porD* (SSO_RS10390) TSS (= DC178 in (7)) |
| WM152<br>WM113 | CGCAACTGTTGGATGAAAGC<br>TCCTCCGCCGATTAATAGGG | F<br>R | Amplification of *b129-a291-c124* region as template for seq reactions in PE *a291* sRNA |
| WM169<br>WM170 | CGAAGATTGCATCACTATACCTTGC<br>CTGAGGAATATGCTTCTCTTACTGC | F<br>R | Amplification of SSO_RS10385 (*permease*)-SSO_RS10390 (*porD*) region template for seq reactions in PE *porD* positive control |
| WM011<br><br><br><br>WM012 | AGCCCTAGACCATGTTTGCAAAAAG<br>CATAATATCTGCGTTAGTTAGGCTC<br>TTTTTAAAGTCTACCTTCTTTTTCG<br>CTTACAATGAGGAAGTCCCTTCTAG<br>CTAGAAGGGACTTCCTCATTGTAAG<br>CGAAAAAGAAGGTAGACTTTAAAAA<br>GAGCCTAACTAACGCAGATATTATG<br>CTTTTTGCAAACATGGTCTAGGGCT | F<br><br><br><br>R | Oligo’s covering whole pT3 promoter region, hybridized to dsDNA, [γ-$^{32}$P]ATP- labelled and used as loading control in PE |
| WM154<br>WM155 | CGATGGCATTGAATTCTGGCGGT<br>TCCAAACCCTTCTGCGTTAAGGC | F<br>R | Targeting N-terminal end of *a291* where antisense sRNA is located, for RT-qPCR |
| WM171<br>WM172 | AATAGAAGGAAACCCGGCCA<br>AACCACACTACCGGATACCC | F<br>R | Targeting C-terminal region of *a291* where only sense RNA is transcribed, for RT-qPCR |
| WM158<br>WM159 | ACTAACGCCACACTCCTGAA<br>GCACCCACTTCATATCACTCC | F<br>R | Targeting *vp1-vp3* region in PCR on cDNA and DNA samples to check for recombination |
